# Small scale variability in soil moisture drives infection of vulnerable juniper populations by invasive forest pathogen

**DOI:** 10.1101/2020.06.09.141739

**Authors:** Flora Donald, Sarah Green, Kate Searle, Nik J. Cunniffe, Bethan V. Purse

## Abstract

The oomycete plant pathogen, *Phytophthora austrocedri*, is an aggressive killer of cypress trees causing severe mortality of Chilean cedar (*Austrocedrus chilensis*) in Argentina since the 1940s and now common juniper (*Juniperus communis* s.l.) in the UK. Rapid mortality of key UK juniper populations was first observed in the early 2000s; the causal agent of mortality was confirmed as *P. austrocedri* in 2012 and the pathogen has now been widely detected - but is not ubiquitous - in juniper populations across Scotland and England. Although juniper has a broad distribution across the northern hemisphere, the UK incidence of *P. austrocedri* remains the only confirmed infection of juniper populations globally. Juniper is an important species for biodiversity, so it is imperative to understand the abiotic and biotic drivers of emergent *P. austrocedri* infection to inform detection, containment and conservation strategies to manage juniper populations across the full extent of its range.

As management of UK juniper populations is primarily conducted at a local level, we investigated field scale drivers of disease – in three, geographically separate populations with different infection histories. Variation in the proportion of juniper showing symptoms - discoloured or dead foliage – was measured using stratified sampling across along key environmental gradients within each 100-hectare population, including juniper density identified from aerial imagery. Potential predictors of infection included altitude, slope, distance to nearest watercourse, soil moisture (mean percentage volumetric water content), area of red deer browsing damage and area of commonly associated vascular plant species. We assessed support in the data for alternative models explaining the spatial distribution of *P. austrocedri* symptoms using full subset covariate selection and Deviance Information Criteria (DIC). Despite differences in environmental gradients and infection histories between populations, area of juniper symptomatic for *P. austrocedri* increased with waterlogging, increasing with soil moisture in sites where soils had higher peat or clay contents, and decreasing with proximity to watercourses where sites had shallower, sandier soils. These results are consistent with key drivers identified at both local and landscape scale in Chilean cedar. Our approach enables identification of site-specific disease management strategies including prioritisation of inspections in microsites with high soil moisture and promoting conservation measures such as creation of sites for natural regeneration in drier microsites to minimise pathogen spread and maximise the resilience of existing juniper populations.

## 1.0 Introduction

The frequency of plant pathogen introductions outside their native ranges is increasing as global trade networks expand (Chapman et al., 2017). Successful establishment of pathogens in these new environments is increasingly being facilitated by degradation of the receiving communities through habitat fragmentation, species turnover and land use change (Chapman et al., 2016; Meentemeyer et al., 2011). Economic losses from plant diseases in the natural environment can result directly from drastic reductions in the extent and viability of host species, increased cost of detection and containment measures, or from indirect losses such as destablisiation of ecosystem functioning from loss of biodiversity or negative visual impacts detering tourists, driving down house prices and increasing local crime rates (Boyd et al., 2013; Kovacs et al., 2011; Mills et al., 2011; Troy et al., 2012). While it is appealing to act immediately to try to control disease outbreaks, the effectiveness of these actions improves as more information on the processes governing spread becomes available, often relying on information that does not exist prior to pathogen introduction (Thompson et al., 2018). Understanding the subset of abiotic and biotic conditions in the invaded range under which introduced pathogens are likely to infect susceptible host populations can improve targeting of such interventions and highlight risk factors for outbreaks in uninvaded locations (Cunniffe et al., 2016).

The oomycete genus *Phytophthora* contains many pathogenic species that adversely impact plant health, forestry and agriculture, necessitating expensive, long-term, landscape scale management. Between 1970 and 1989, 11 *Phytophthora* species were introduced to China, 12 to the UK and 16 to the USA (Barwell et al., in review). In the two decades following, the number of additional species introduced at least doubled in China (20) and the UK (29) and increased five-fold in the USA (54). While not all of these species established, some of them have caused serious tree mortality with dramatic landscape and economic consequences. In Western Australia 282 000 ha of Jarrah (*Eucalyptus marinata*) have been lost to *P. cinnamomi* (Boyd et al., 2013), while trade of Port Orford cedar (*Chamaecyparis lawsoniana*) in the north-western United States was almost eliminated by *P. lateralis* (Hansen, 2015). Meanwhile, millions of coast live oak (*Quercus agrifolia*) and tanoak (*Notholithocarpus densiflorus*) trees in California and Oregon, and 18 000 ha of Japanese larch (*Larix kaempferi*) in the UK and Ireland have been killed by *P. ramorum* (Meentemeyer et al., 2011; O’Hanlon et al., 2018; Peterson et al., 2014).

First described in 2007, *Phytophthora austrocedri* Gresl. & E. M. Hansen has caused widespread mortality of Chilean cedar (*Austrocedrus chilensis*) in Argentina since the 1940s (Greslebin et al., 2007; Greslebin and Hansen, 2010). The pathogen is homothallic and is potentially spread by both asexual, motile zoospores dispersed through any form of moving water, and sexually produced, thick-walled oospores that can remain viable for extended periods of time and be translocated in soil (Green et al., 2015; Henricot et al., 2017a). Infection usually starts in the roots before spreading into the cambium and phloem, creating necrotic lesions that can extend to the full width of each layer, starving whole branches, trunks or trees of water and nutrients causing rapid defoliation and mortality (Green et al., 2015).

Symptoms were first brought to the attention of UK plant pathologists in the mid-2000s, when significant numbers of symptomatic juniper (*Juniperus communis* L. s. l.) could be observed in two of the larger populations (Glenartney and Haweswater), but *P. austrocedri* was not confirmed as the causal agent of mortality until 2012 following isolation and confirmation of Koch’s postulates (Green et al., 2012). In the UK the pathogen is present as a single genetic lineage exhibiting no diversity in nucleic and mitochondrial loci, suggesting introduction and spread of a single clonal strain (Henricot et al., 2017a). The extent of juniper decline varies, with some populations showing wholesale dieback of bushes compared to others with only localised patches of symptoms, suggesting populations have been infected at different times and/or different site conditions promote different rates of spread.

Although interceptions of infected cypress and juniper trees in Scotland, England and Germany confirm that infected material is being traded (Green et al., 2015; Werres et al., 2014) the outbreak in British juniper populations in the wider environment remains the only detected infection of a natural host population outside Argentina (Green et al., 2012). Globally, juniper has a large, circumboreal distribution extending across the northern hemisphere but as no investigation of environmental drivers of infection has been undertaken, the proportion of juniper vulnerable to *P. austrocedri* infection is unclear.

In the UK, juniper has a wide but discontinuous distribution occupying much of Scotland and northern England and remaining as scattered populations in southern England, Wales and Northern Ireland (Fig. 1). Populations are undergoing long term declines in most areas as a result of burning, afforestation, over-grazing, under-grazing, increased levels of diffuse pollution (particularly nitrogen) and poor germination following warmer winter temperatures (Broome and Holl, 2017; Clifton et al., 1997; Long and Williams, 2007; Sullivan, 2003; Verheyen et al., 2009; Walker et al., 2017; Ward and Shellswell, 2017). It is difficult to estimate the area of juniper lost specifically to disease but infected populations are widespread across Scotland and England, where juniper occupancy of 10 × 10 km cells reportedly declined by 23 % and 44 % respectively between 2000 and 2016 (Plantlife, 2015).

**Figure 1.**
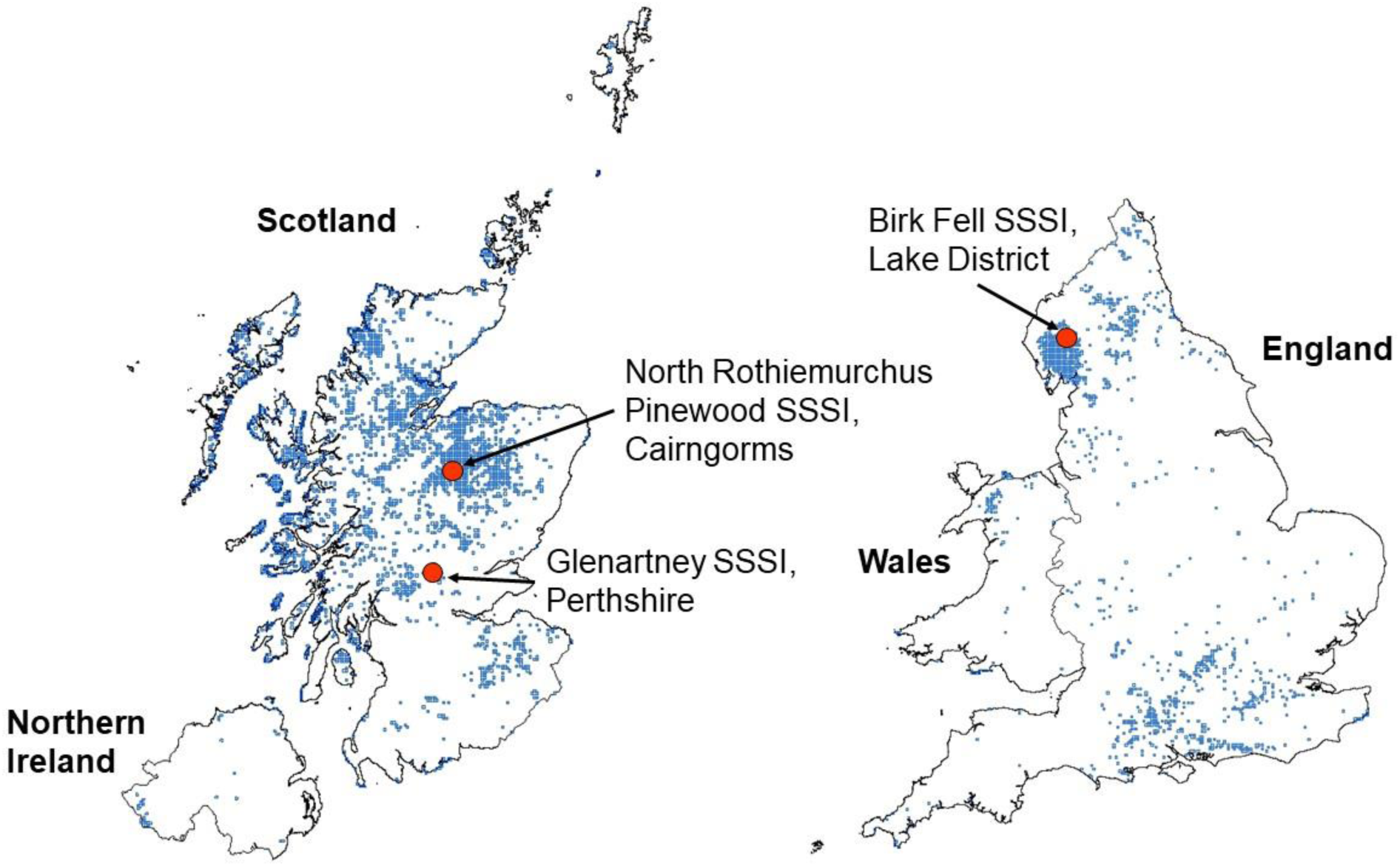
SSSI name and location of the three juniper study populations mapped against the distribution of UK juniper (*Juniperus communis* s. l.) at 2 × 2 km resolution (shown in blue) recorded during the period 2000 – 2017 (Botanical Society of Britain and Ireland, 2017).

The societal and environmental value of woodlands was recognised in the 2014 Tree Health Management Plan for England as several times higher than the commercial value of forestry (Department for Environment Food & Rural Affairs, 2014a) and the Scottish Plant Health strategy identifies plant health in the natural environment as integral to the £1.8 billion rural economy (The Scottish Government, 2016). Loss of UK juniper populations to *P. austrocedri* could be significant as the species is highly ecologically important as a dominant component of many habitats including woodland, scrub, heath, dune and calcareous grassland, as a nurse species ameliorating environmental conditions and protecting other seedlings from herbivory, a rare source of winter food and nesting habitat for birds, and a host of many specialist fungi and insects (Thomas et al., 2007; Ward and Shellswell, 2017; Wilkins and Duckworth, 2011).

As juniper occupies such a broad variety of habitats, trees are subject to different environmental conditions and land uses that may alter their susceptibility to disease. Epidemics occur across a range of spatial scales, arising first as microscopic infections that can spread to whole plants, populations and landscapes (Gilligan and Van Den Bosch, 2008). Successful disease control requires matching of the scale of management to the inherent scale of spread as mediated by host population connectivity and pathogen dispersal distances (Cunniffe et al., 2016). Transmission of soil borne pathogens is likely to occur across short distances resulting in highly aggregated infection prevalence and spatially variable exposure to pathogens within host populations (Penczykowski et al., 2018). However, very few studies of soil borne Phytophthoras investigate spread at field scale and those that do use small (< 20) sample sizes (La Manna and Matteucci, 2012; La Manna and Rajchenberg, 2004a; Nagle et al., 2010; Tippett et al., 1989).

We measured juniper symptoms in 147 quadrats across three, geographically separate juniper populations with contrasting infection intensities and analysed the data using Generalised Linear Mixed Models (GLMMs) to compare drivers of spatial variation in symptoms at field scale. Correlative approaches, such as GLMMs, are appropriate tools to perform such exploratory analyses as they can accommodate a broad range of potential covariates as is necessary when drivers of pathogen dispersal and spread in the invaded range are poorly understood (Purse and Rogers, 2009).

We expected *P. austrocedri* infection of juniper would exhibit similar responses to environmental covariates as infected Chilean cedar, which occupies a similarly diverse range of ecotypes. Population level studies in Argentina found area of foliage symptoms increased in microsites situated at low altitude with poor soil drainage, flat slopes, close proximity to watercourses and fine soil textures, with greater infection of female cedars because they typically occupy wetter microsites (Baccalá et al., 1998; V. a El Mujtar et al., 2012; La Manna et al., 2008; L. La Manna and Rajchenberg, 2004; Ludmila La Manna and Rajchenberg, 2004).

In addition, we expected that the area of symptoms in juniper would increase with i) increasing host density, as inoculum production is likely to increase with availability of host tissue and dispersal distances between roots will be reduced (Anderson and May, 1986; Dillon et al., 2014); and ii) increased ungulate herbivory, and proximity to deer and sheep tracks and lie-ups, as evidence for increased exposure to inoculum transported in soil on herbivore hooves (Jules et al., 2002; La Manna et al., 2013).

Given the contrast in abiotic and biotic conditions occupied by each population, we further expected that the relative importance of the investigated covariates would vary between locations and require site specific strategies to manage individual juniper populations.

## 2.0 Methods

### 2.1 Study Areas

Three infected juniper populations from where *P. austrocedri* had previously been isolated (Henricot et al., 2017) were selected to best represent the diversity of climatic, topographic and edaphic conditions occupied by juniper. Two populations are located in Scotland: one in Perthshire (P) and one in the Cairngorms (C), and one population is situated in the Lake District (LD) in the north of England. In all three locations, the juniper population is a component feature of a Special Area of Conservation designated habitat and a qualifying interest of a Site of Special Scientific Interest (SSSI; Fig. 1).

Each population contained 100 – 130 hectares of continuous juniper cover. The greatest area of mortality was observed in the Perthshire population; symptoms were first reported in 2004 and represented a 20% decline in area of live juniper trees compared to a 1983 baseline survey (Tene et al., 2007). The precise duration of infection in the Lake District and Cairngorms populations is unknown, but *P. austrocedri* symptoms were first noted after 2010, and a lower proportion of symptomatic juniper was observed at both sites potentially consistent with a more recent introduction.

### 2.2 Quadrat stratification

Juniper was sampled using 10 × 10 m quadrats from pre-selected locations stratified according to the area and density of juniper, altitude, slope and distance to watercourses. A 2010 distribution map of the Perthshire juniper population derived from 15 cm full colour (RGB) and false colour infrared (CIR) imagery was provided by Scottish Natural Heritage (Whittome, 2010) and distribution maps of the remaining two populations were classified from 25 cm RGB imagery supplied for 2010 by NeXTPerspectives™. The classification methods are described in Appendix A.

To capture differences in juniper abundance and density, the area of juniper predicted by the image classifications was measured in 10 × 10 grid cells using landscape class statistical functions in the SDMTools package (Van der Wal et al., 2014) implemented in R v. 3.4.2 (R Core Team, 2017). To understand if each 10 × 10 m cell was isolated from other juniper stands or part of a larger stand, the area of juniper in 30 × 30 m including each 10 × 10 m grid cell was also calculated, producing distributions of juniper % cover at each scale for each study population (Fig. 2). These distributions were used to devise eight categories to describe juniper abundance that could be easily identified in the field. Each 10 × 10 m cell containing juniper was assigned to one of four categories describing juniper % cover in 10 × 10 m (≤ 10, 11-25, 26-50, > 51 %) and to one of two categories characterising the area of juniper surrounding the 10 × 10 m cell as isolated from (< 20 %), or contiguous with (> 21 %), juniper growing in the wider 30 × 30 m.

**Figure 2.**
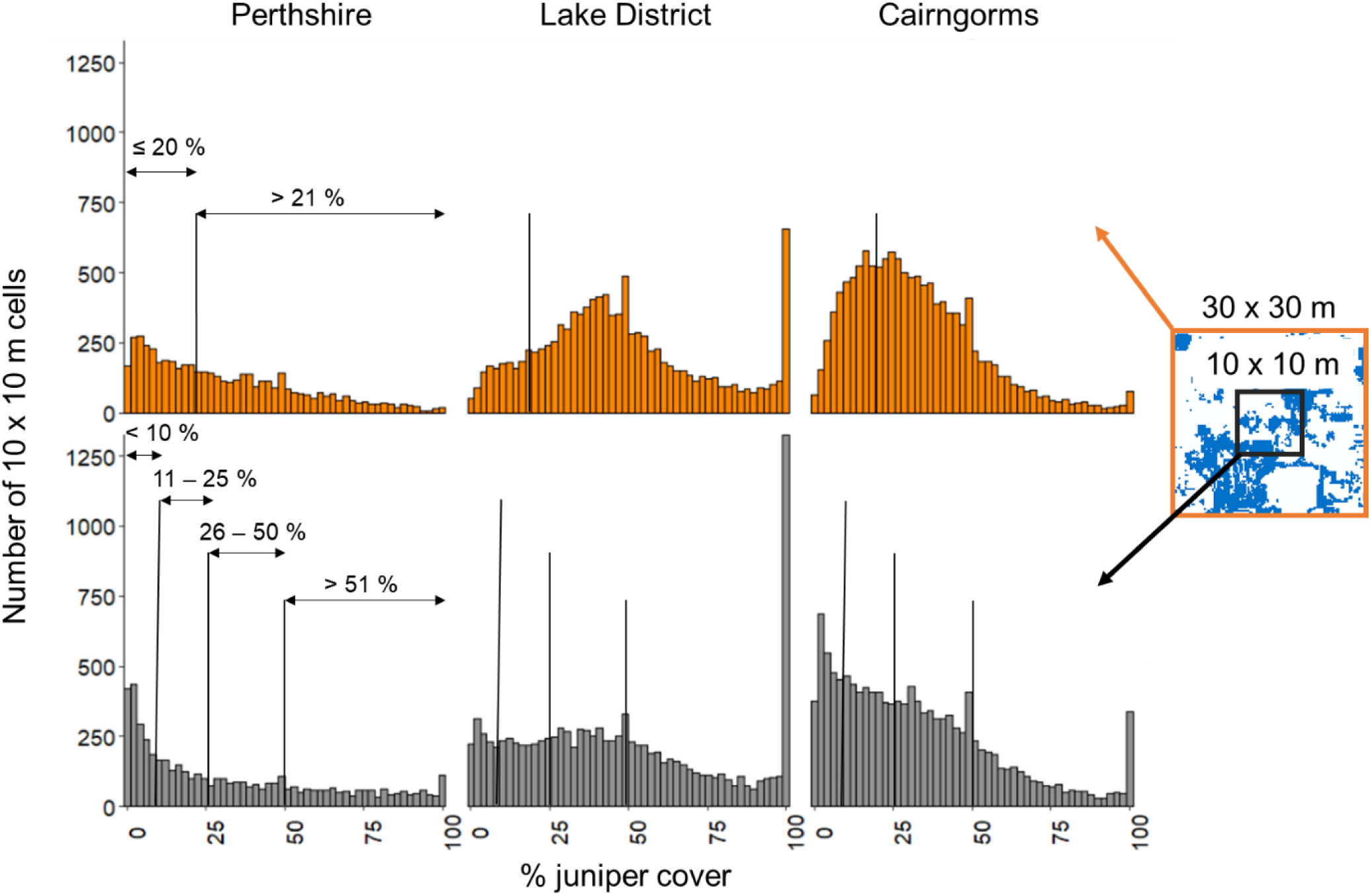
Number of 10 × 10 m cells per study population with estimated % cover of juniper (shown in blue) across 10 × 10 m (grey) cells and the surrounding 30 × 30 m (orange). Thresholds used to divide juniper % cover into categories at each scale are marked with black lines.

Layers of slope and aspect were calculated from the resampled NeXTPerspectives™ 10 m DEM using the *terrain* function in the raster package (Hijmans, 2016). Slope, aspect and altitude were then extracted to the centroid of each 10 × 10 m grid cell containing juniper. Euclidean distance (m) from the nearest watercourse to each 10 m grid cell centroid was measured from a rasterised version of the 50 m digital rivers network (Moore et al., 2000).

Fifty 10 × 10 m cells per study population were randomly selected for sampling in proportion to the total number of cells assigned to each abundance category. This was repeated five times for each population. After each run, selected cells were plotted across the altitudinal, slope and watercourse proximity gradients occupied by juniper at each location and the selection that captured the widest distribution of samples along each of the three gradients was chosen as sampling locations.

### 2.3 Survey of spatial patterns in juniper symptoms

Quadrat sampling was carried out over five days at each location in October 2017. Quadrats were geo-located using ArcPad v. 10.2 on a Panasonic FZ-GI tablet with GPS accuracy to 3 m. To minimise transference of inoculum across populations, areas of high and low infection were visited on different days and all equipment breaking the soil surface (e.g. marker poles, soil moisture probes) was disinfected between quadrats. All other equipment was thoroughly disinfected between study populations.

Juniper quadrats were placed as close to pre-selected locations as was possible to meet the abundance criteria by estimating the area of juniper in 10 × 10 m and scoring abundance in 30 × 30 m as a binary measure of more or less than 20 %. The area of symptomatic juniper was measured as a fraction of the total area of juniper present in each quadrat, where symptoms constituted foliage discolouration and dead needles (retained or dropped) that extended to a minimum of a whole branch and did not result from either browsing or mechanical damage. Where a distinctive phloem lesion typical of *P. austrocedri* could be found, a 500 mg tissue sample was collected from one representative symptomatic tree per quadrat. The sampled tissue was stored at - 20 °C until quantitative real-time PCR (qPCR) could be carried out following the protocol described in Mulholland et al. (2013) to verify the consistent presence of *P. austrocedri* across each population.

### 2.4 Abiotic and biotic predictors

We measured a suite of potential abiotic and biotic predictors of spatial patterns in *P. austrocedri* symptoms and included these in statistical models for each population (Table 1). The following predictors were measured in each field quadrat. The binary observation of ≤ 20 % or > 21 % juniper cover across 30 × 30 m to distinguish between quadrats situated in isolated or contiguous juniper stands was included in the model as juniper “density”. The area of juniper bearing berries was used to estimate the area of female juniper. Area of herbivore damage was measured as the area of bark stripping plus any resulting dead branches / stems (i.e. mechanical breakage from wind or snow damage was excluded). This metric was not included in Cairngorm models as herbivory was only detected in nine quadrats, encompassing an area greater than 10 cm^2^ in only three quadrats.

**Table 1.** List of covariates included in full subset model selection for each population (P = Perthshire, LD = Lake District, C = Cairngorms). Number of sampled quadrats: P = 51, LD = 46, C = 50.

| Measurement category | Specific measurement | Population |  |  |
| --- | --- | --- | --- | --- |
|  |  | P | LD | C |
| Juniper density<br>30 x 30 m quadrat | ≤ 20 % or > 21 % juniper cover | x | x | x |
| Juniper metrics<br>10 x 10 m quadrat | Area of juniper bearing berries (m<sup>2</sup>) | x | x | x |
|  | Area of herbivore damage (m<sup>2</sup>) | x | x |  |
| Soil moisture | Mean of point samples across quadrat (% VWC) | x | x | x |
| Vascular plant indicators | Area of individual target taxa (m<sup>2</sup>) | x | x | x |
| Watercourse proximity | Euclidean distance from quadrat centroid to nearest mapped river (m) | x | x | x |
| Grazing activity | Distance from quadrat centroid to nearest deer or sheep track (m) | x |  |  |
| Topographic metrics<br>extracted to 10 x 10 m<br>quadrat centroid | Altitude (m) | x | x | x |
|  | Slope (°) | x | x | x |
|  | Aspect (°) |  |  | x |
| Soil type (250 m<br>resolution) extracted to<br>10 x 10 m quadrat<br>centroid | **Perthshire:**<br>Brown forest Balrownie<br>Brown forest Gourdie<br>Organic soil, peaty gleys<br>Non-calcareous gleys<br>Peaty gleys Balrownie | x |  |  |
|  | **Cairngorms:**<br>Humus-iron podzols; some brown forest soils,<br>noncalcareous gleys and peaty gleys<br>Humus-iron podzols; some peaty gleys and humic gleys |  |  | x |
| Habitat<br>(NVC community) | **Perthshire:**<br>W19 *Juniperus communis* woodland<br>Acid grasslands (U4, U20, U24)<br>Mires (M10, M23)<br>Mosaic (U5, M15) | x |  |  |
|  | **Cairngorms:**<br>H12 *Calluna vulgaris* – *Vaccinium myrtillus* heath<br>W18 Scots pine woodland with heather<br>U4 *Festuca ovina* - *Agrostis capillaris* - *Galium saxatile*<br>grassland<br>Coniferous plantation |  |  | x |

Soil moisture was measured as % volumetric water content (VWC) using a FieldScout TDR 300 probe. Shallow soil and surface rock only permitted measurements using the 3.8 cm depth setting across the Lake District population, whereas measurements were collected at 20 cm depth in Perthshire and the Cairngorms. Measurements were collected from i) areas within each quadrat where juniper was absent, ii) under asymptomatic juniper and iii) under symptomatic juniper. An equal number of measurements (minimum four) was collected from each category present, resulting in eight to twelve point sample measurements from which mean soil moisture was calculated (% VWC).

Area of vascular plant taxa present in each quadrat was recorded according to a target list (Appendix B) of taxa chosen to indicate placement of microsites along soil moisture, nitrogen and pH gradients.

Mapping in the field was carried out using the tracking function in ARCpad, to record any watercourses additional to the 50 m digital rivers network (Moore et al., 2000). These were merged with the original dataset and used to recalculate the watercourse proximity (m) metric for each quadrat. Clearly visible deer and sheep tracks were also mapped and proximity to sampled quadrat centroids measured as an alternative way to measure the risk of inoculum transference to juniper from herbivores, but stocking density and ground condition only permitted collection of a reliable dataset from Perthshire.

The remaining covariates were obtained from existing GIS datasets. Altitude, slope and aspect were extracted to each quadrat centroid from the resampled NeXTPerspectives™ 10 m layers prepared for the plot stratification. Aspect was not included in the Perthshire or Lake District models as more than 60 % of quadrats at each location were clustered in the same octant.

The soil type underlying each quadrat centroid was extracted from 250 m resolution datasets, obtained from a digitised version of the soil map produced by Forbes (1984), the Soilscapes dataset (Farewell et al., 2011) and the National Soil Map of Scotland (James Hutton Institute, 2011) for the Perthshire, Lake District and Cairngorms populations respectively. Soil type was omitted from model selection for the Lake District population as at 250 m resolution all of the quadrats were placed in “freely draining acid loamy soils over rock” (Farewell et al., 2011).

To test if a broader description of the vegetative community is a better predictor of *P. austrocedri* symptoms, because it captures more information about edaphic conditions than the presence of individual taxa, National Vegetation Classification (NVC) community data, supplied by Scottish Natural Heritage (2017), was included as a covariate for the Perthshire and Cairngorms populations (Table 1). The eight Perthshire communities were simplified to these four broad types (Table 1), amalgamated as follows: acid grasslands (U4 *Festuca ovina* - *Agrostis capillaris* - *Galium saxatile*; U20 *Pteridium aquilinum* - *Galium saxatile*; U24 *Arrhenatherum elatius* - *Geranium robertianum*), mires (M10 *Carex dioica* – *Pinguicula vulgaris*; M23 *Juncus effusus/acutiflorus* – *Galium palustre*) and mosaic communities suggesting transition from drier to wetter soil (U5 *Nardus stricta* – *Galium saxatile*; M15 *Trichophorum germanicum* - *Erica tetralix* wet heath) (Rodwell, 1991). No NVC data was released for the Lake District population.

### 2.5 Model specification

To investigate the relationships between the area of *P. austrocedri* symptoms and environmental covariates, we used a Bayesian beta-binomial Generalised Linear Mixed Model (GLMM) fitted using the Integrated Nested Laplace Approximation (INLA) method with the R-INLA package (Rue et al., 2009) implemented in R v. 3.4.2 (R Core Team, 2017).

Models were fitted to the number of square metres of symptomatic juniper in each 10 × 10 m quadrat. Using the beta-binomial distribution enabled us to take account of the area of juniper in each cell while allowing the probability of infection to have extra variation associate with spatial clustering of symptoms (overdispersion), thereby accounting for the high frequency of quadrats that contain wholly asymptomatic or symptomatic juniper (Hughes and Madden, 1993).

In particular, our model used the *q* = 12 environmental covariates for *i*^th^ the location {*x*_*j,i*_|1 ≤ *j* ≤ 16} to estimate the mean probability of infection via a logit link function

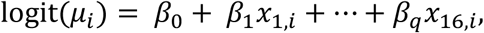

in which β_0_ is an intercept and β_j_ is the regression coefficient for the *j*^th^ predictor. This estimate of the mean probability is then used to predict the area of symptomatic juniper in the *i*^th^ location (*η*_*i*_) via

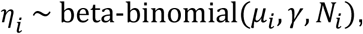

in which *N*_i_ is the total area of juniper in the *i*^th^ cell (m^2^) and *γ* is the overdispersion parameter of the beta-binomial distribution (that was assumed to be constant across all cells at each site). In our Bayesian estimation procedure all regression parameters, including the intercept, were assumed to have minimally informative priors of the form

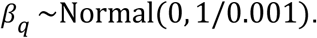

### 2.6 Model selection

All covariates were centred and standardized prior to model fitting and no pairs of covariates used in any models were correlated with a Pearson r^2^ value ≥ 0.6 (Appendix D). Models were run in two stages. We first performed a full subset selection using all possible combinations of covariates marked against each population (Table 1) except the vascular plant indicators, producing 1023 models for both Perthshire and the Cairngorms, and 127 models for the Lake District, which had seven as opposed to ten covariates.

Model fit was compared using the Deviance Information Criterion (DIC), a Bayesian generalisation of the Akaike Information Criterion (AIC) derived as the mean deviance adjusted for the estimated number of parameters in the model to provide a measure of out-of-sample predictive error (Gelman and Hill, 2006). The model with the lowest DIC is the model with the most support in the data, but the set of models with DICs within two units of the top model DIC are considered to have equivalent support in the data and formed the “top model set”.

The area of each vascular plant indicator, present in ten or more quadrats at each population, were then added to the formulae for the top model set per population to assess (using DIC) if the addition of any one indicator improved model fit. Nine additional models were run for Perthshire and the Lake District, and ten for the Cairngorms (Table 3).

**Table 2.** Comparison of surveyed juniper population covariate means ± 1 s.d., and ranges, measured from 10 × 10 m quadrats. P = Perthshire, LD = Lake District and C = Cairngorms study populations. Only numerical / binary coded variables included in the first stage of GLMM modelling (i.e. excluding species indicators) are displayed. Covariates not included in models for specific populations are greyed out.

| Covariate | Mean ± s.d.per quadrat | | | Range per quadrat | | |
| --- | --- | --- | --- | --- | --- | --- |
|  | P | LD | C | P | LD | C |
| Area of symptomatic juniper (m<sup>2</sup>) | 19 ± 20 | 13 ± 18 | 7 ± 11 | 0 - 79 | 0 - 70 | 0 - 45 |
| Area of juniper (m<sup>2</sup>) | 34 ± 28 | 44 ± 27 | 33 ± 22 | 2 - 99 | 1 - 90 | 1 - 80 |
| Juniper density<br>(% quadrats nested in<br>30 x 30 m with > 21<br>% juniper cover) | 35 | 72 | 70 | NA | NA | NA |
| Area of juniper bearing berries (m<sup>2</sup>) | 10 ± 12 | 9 ± 15 | 4 ± 9 | 0 - 49 | 0 - 66 | 0 - 49 |
| Area of herbivore damage (m<sup>2</sup>) | 2 ± 8 | 0 ± 1 | 0 ± 1 | 0 - 50 | 0 - 3 | 0 - 7 |
| Mean soil moisture<br>(% VWC) | 45 ± 12 | 63 ± 17 | 33 ± 9 | 28 - 84 | 33 - 108 | 15 - 52 |
| Watercourse proximity<br>(m) | 42 ± 37 | 144 ± 209 | 240 ± 193 | 1 - 174 | 0 - 840 | 2 - 820 |
| Grazing activity (m) | 16 ± 16 | 72 ± 68 | 31 ± 49 | 0 - 70 | 1 - 212 | 1 - 213 |
| Altitude (m) | 250 ± 30 | 248 ± 63 | 303 ± 21 | 182 - 314 | 150 - 384 | 256 - 334 |
| Slope (°) | 11 ± 4 | 32 ± 7 | 6 ± 4 | 5 - 23 | 15 - 45 | 1 - 20 |
| Aspect (°) | 158 ± 145 | 276 ± 53 | 174 ± 99 | 2 - 357 | 88 - 342 | 10 - 348 |

**Table 3.**
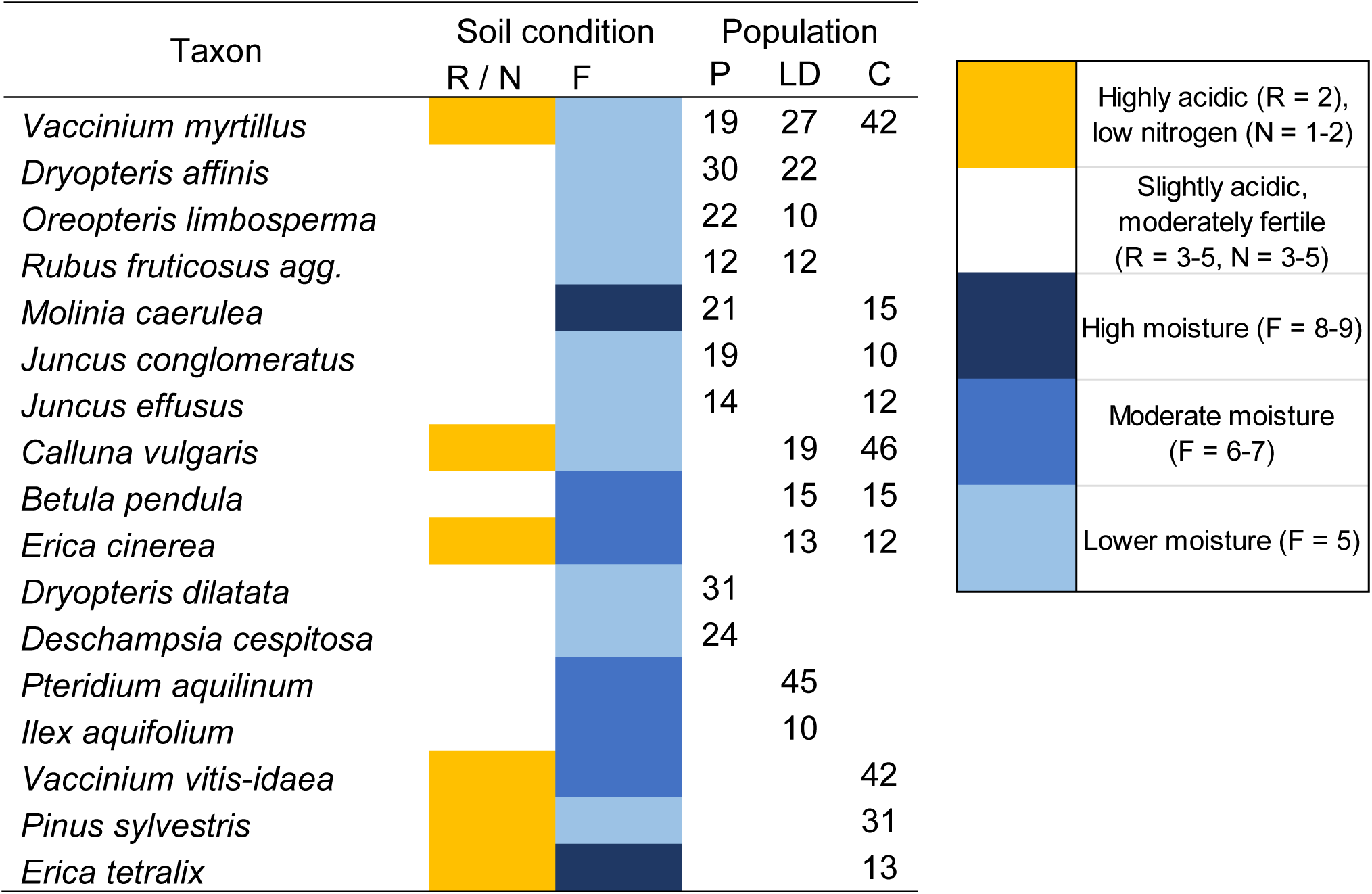
Number of 10 × 10 m quadrats containing vascular plant indicators (where present in ≥ 10 quadrats). P = Perthshire, LD = Lake District, C = Cairngorms juniper populations. Taxa are colour coded according to soil condition categories devised from Ellenberg reaction (R), nitrogen (N) and moisture (F) values given in Hill, Preston, & Roy (2004).

| Taxon | Soil condition |  | Population |  |  |
| --- | --- | --- | --- | --- | --- |
|  | R / N | F | P | LD | C |
| *Vaccinium myrtillus* | Yellow | Light blue | 19 | 27 | 42 |
| *Dryopteris affinis* |  | Light blue | 30 | 22 |  |
| *Oreopteris limbosperma* |  | Light blue | 22 | 10 |  |
| *Rubus fruticosus* agg. |  | Light blue | 12 | 12 |  |
| *Molinia caerulea* |  | Dark blue | 21 |  | 15 |
| *Juncus conglomeratus* |  | Light blue | 19 |  | 10 |
| *Juncus effusus* |  | Light blue | 14 |  | 12 |
| *Calluna vulgaris* | Yellow | Light blue |  | 19 | 46 |
| *Betula pendula* |  | Blue |  | 15 | 15 |
| *Erica cinerea* | Yellow | Blue |  | 13 | 12 |
| *Dryopteris dilatata* |  | Light blue | 31 |  |  |
| *Deschampsia cespitosa* |  | Light blue | 24 |  |  |
| *Pteridium aquilinum* |  | Blue |  | 45 |  |
| *Ilex aquifolium* |  | Blue |  | 10 |  |
| *Vaccinium vitis-idaea* | Yellow | Blue |  |  | 42 |
| *Pinus sylvestris* | Yellow | Light blue |  |  | 31 |
| *Erica tetralix* | Yellow | Dark blue |  |  | 13 |

|  |  |
| --- | --- |
| Yellow | Highly acidic (R = 2),<br>low nitrogen (N = 1-2) |
| White | Slightly acidic,<br>moderately fertile<br>(R = 3-5, N = 3-5) |
| Dark blue | High moisture (F = 8-9) |
| Blue | Moderate moisture<br>(F = 6-7) |
| Light blue | Lower moisture (F = 5) |

To assess the importance of covariate effects, we summarised marginal posterior distributions using 95 % (0.025 and 0.975 quantiles of the posterior distribution) Bayesian credible intervals (BCI). The relationship between each covariate and the area of symptoms is considered strongest where BCI do not bridge zero, very strong when ≥ 0.95, strong when ≥ 0.90 – 0.94, and weak when ≥ 0.80 – 0.89 of the BCI are above or below zero. Where a strong or very strong relationship was found between the area of symptoms and a covariate, we report the percentage of the posterior predicted data that overlaps zero as calculated in R v. 3.5.2 (R Core Team, 2018) using the *rollmean* function in the zoo package (Zeileis and Grothendieck, 2005).

Model validation was performed using root mean-square error (RMSE) calculated between the predicted posterior mean values and the corresponding mean sampled area of symptomatic juniper. The residuals of the top models were checked for spatial autocorrelation using Moran’s I statistic implemented using the *correlog* function in R package ncf v 1.2.8 (Bjornstad, 2019). Pairs of plots were divided into different distance bins at 100 m intervals between 0 and the maximum distance between plot pairs for each site and the Moran’s I value was then calculated for each distance bin. One hundred paired distances were randomly resampled per distance bin to assess Moran’s I correlation significance (Appendix D).

## 3.0 Results

### 3.1 Prevalence of symptoms of *P. austrocedri* infection

Fifty-five percent of juniper surveyed in the Perthshire population showed symptoms compared to 28 % in the Lake District and 23 % in the Cairngorms populations, consistent with a possible earlier pathogen introduction in Perthshire (Fig. 3). Though quadrats containing no symptomatic juniper were found in all three populations, the mean area of symptomatic juniper found in Perthshire quadrats was 19 ± 20 m^2^ out of a mean 34 ± 28 m^2^ area of juniper, compared to a mean of 7 ± 11 m^2^ of symptomatic juniper in quadrats from the Cairngorms population where the mean juniper cover found per quadrat was similar (33 ± 22 m^2^). The mean area of juniper in the Lake District quadrats was higher (44 ± 27 m^2^) with an intermediate mean area of symptomatic juniper (13 ± 18 m^2^).

**Figure 3.**
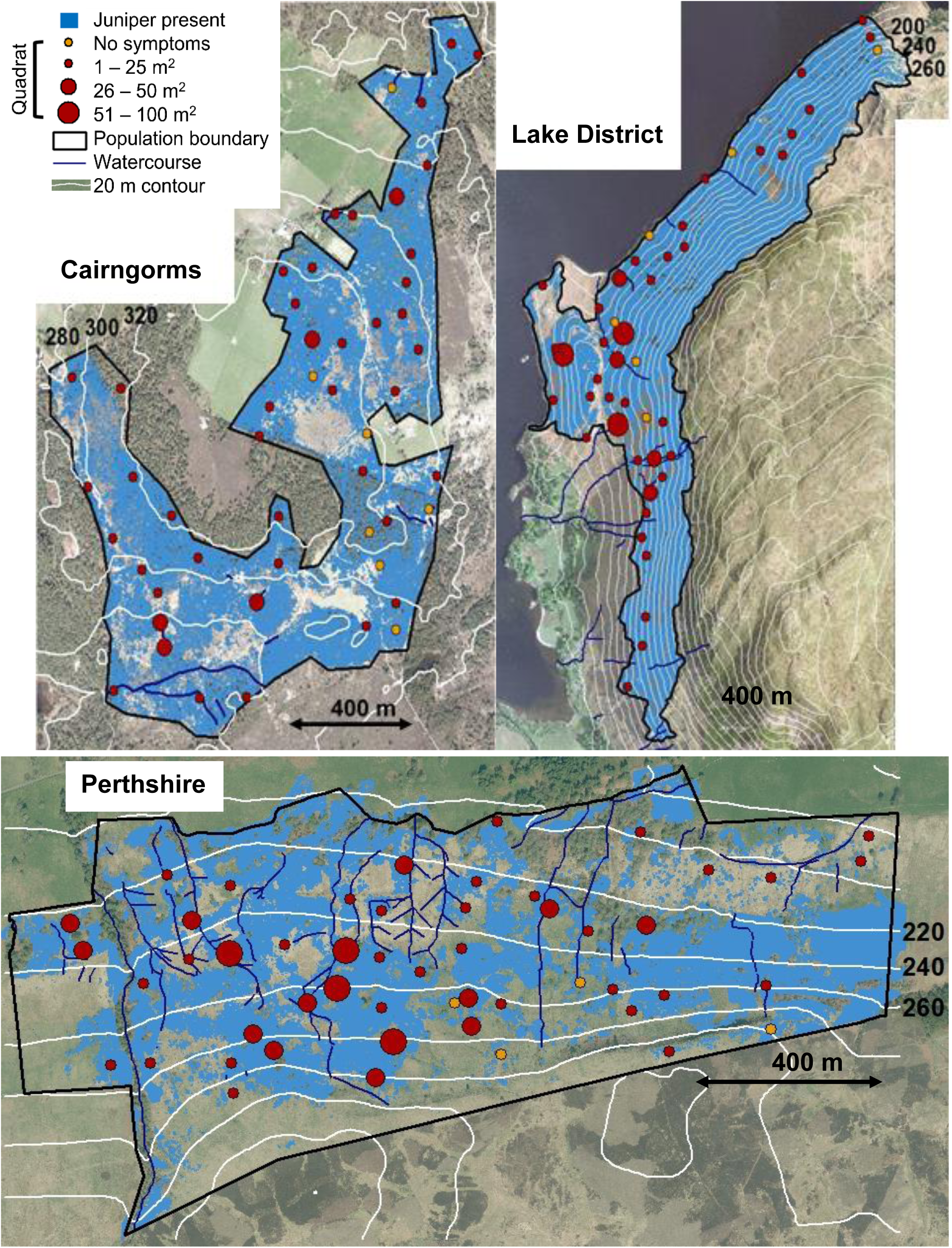
Map of surveyed juniper populations showing the distribution of juniper in relation to the watercourse, altitude and slope covariates used to stratify sampling. The distribution of samples, collected in 10 × 10 m quadrats, is shown with circles coloured orange where no *P. austrocedri* symptoms were found and red where symptoms were present. Circle size corresponds to categories representing the area of symptoms estimated in each quadrat. Imagery licensed to UK Centre for Ecology & Hydrology for PGA through Next Perspectives™

Because detection of symptomatic lesions is limited to above-ground live tissue, qPCR results are less reliable indicators of infection than symptoms. However, positive qPCR results were obtained across the full extent of each population giving confidence that site-wide observations of symptoms result from *P. austrocedri* infection (Appendix C).

The mean, standard deviation and range of covariates measured and tested in models across all three populations is shown in Table 2. The Perthshire population is characterised by fragmented juniper stands, only 35 % of quadrats contained more than 21 % juniper cover across 30 × 30 m compared to c. 70 % in the Lake District and Cairngorm populations where juniper grows in denser stands (Fig. 3, Table 2). Perthshire population quadrats were never further than 174 m from a river or drain, compared to 840 and 820 m in the Lake District and Cairngorms populations respectively (Fig. 3, Table 2). The Lake District population occupied the largest range of altitude (234 m compared to 132 m in Perthshire and 78 m in the Cairngorms) with up to 45° slopes compared to just 20° in both Scottish populations (Fig. 3; Table 2). The Cairngorms population had the driest soil conditions across the quadrats (Table 2) with mean soil moisture of 33 % VWC, which is 27 % and 48 % drier than the mean soil moisture found across Lake District and Perthshire quadrats respectively.

Nine vascular plant indicators were present in ≥ 10 quadrats in the Perthshire population, nine in the Lake District and ten in the Cairngorms (Table 3). The mix of indicators recorded highlights the difference in microsites occupied by the juniper study populations (Table 3). Of 42 target indicators, only one, V*accinium myrtillus*, was present at all three study populations while quadrat frequency for the remaining indicators varied from 19 – 42 quadrats. Indicators of drier, moderately fertile soils were only present in the Lake District quadrats, where no indicators of high soil moisture and three indicators of highly acidic, infertile microsites were also found. In contrast, there were two indicators of high soil moisture in the Cairngorms, six for highly acidic, infertile soils (four present in > 40 of 50 quadrats) and no indicators of drier, moderately fertile microsites (Table 3). Quadrats from Perthshire were dominated (both in terms of species composition and prevalence across quadrats) by seven taxa indicating moderate soil moisture (Table 3). However, although only one taxon (*Molinia caerulea*) indicating high soil moisture was recorded, it was found in 21 of the 51 quadrats, suggesting widespread, continuous waterlogging across the site.

### 3.2 Abiotic and biotic drivers of spatial variability in disease symptoms of *P. austrocedri*

The full subset selection modelling resulted in one top model each containing abiotic and biotic covariates for the Perthshire and Lake District populations, and two models for the Cairngorms population (Table 4). All models included a strong relationship between increasing area of *P. austrocedri* symptoms and a measure of increasing soil moisture (Table 5). When the area of different vascular plant indicators was added to these models, this resulted in one top model with improved fit for each population, with strong, positive relationships between increasing symptoms and increasing soil moisture still included but additionally identifying taxa that aid identification of microsites vulnerable to *P. austrocedri* infection in different habitats (Table 4). Across all sites, models with abiotic and biotic covariates vastly outperformed the null model with no covariates.

**Table 4.** Model results (DIC, RMSE, dispersion and list of covariates present) for each surveyed population, comparing the null model with the top set of models produced before and after addition of vascular plant indicators.

| Juniper population | Model | DIC | RMSE | Dispersion | Covariates |
| --- | --- | --- | --- | --- | --- |
| Perthshire | Without indicators | 167.45 | 43.89 | 0.48 | Juniper density, berry bearing, herbivore damage, soil moisture, watercourse proximity, grazing activity, altitude, slope, soil type, habitat |
|  | With indicators | 134.96 | 42.59 | 0.45 | As above with area of *Dryopteris dilatata* |
|  | Null | 301.98 | 37.04 | 0.55 | N/A |
| Lake District | Without indicators | 264.64 | 32.64 | 0.44 | Juniper density, berry bearing, soil moisture, watercourse proximity, altitude, slope |
|  | With indicators | 208.84 | 22.99 | 0.42 | As above with area of *Rubus fruticosus* agg. |
|  | Null | 301.77 | 41.54 | 0.57 | N/A |
| Cairngorms | Without indicators | 226.71 | 21.22 | 0.33 | Juniper density, berry bearing, soil moisture, aspect, soil type, habitat |
|  |  | 225.40 | 21.38 | 0.33 | Juniper density, berry bearing, soil moisture, slope, aspect, soil type, habitat |
|  | With indicators | 191.92 | 21.09 | 0.30 | As above with area of *Erica tetralix* |
|  | Null | 293.63 | 43.92 | 0.58 | N/A |

**Table 5.** Posterior estimates (mean, standard deviation (SD), 2.5 % and 97.5 % quantiles, and % that does not bridge zero) for fixed effects included in the top model set for the Perthshire juniper population.

|  | Covariate | Mean | SD | 2.5% | 97.5% | % does not bridge zero |
| --- | --- | --- | --- | --- | --- | --- |
| *Without indicator species (DIC = 167.45)* |  |  |  |  |  |  |
|  | Soil moisture | 0.59 | 0.31 | 0.01 | 1.21 | 100.00 |
|  | Altitude | -0.69 | 0.43 | -1.57 | 0.13 | 92.71 |
|  | Herbivore damage | -0.39 | 0.29 | -1.01 | 0.14 | 90.20 |
|  | Juniper density | 0.29 | 0.25 | -0.20 | 0.77 | 87.70 |
|  | Intercept | 0.05 | 21.22 | -41.61 | 41.67 | 50.17 |
|  | Berry bearing | 0.21 | 0.25 | -0.27 | 0.70 | 80.19 |
|  | Grazing activity | -0.17 | 0.26 | -0.67 | 0.34 | 75.21 |
|  | Watercourse proximity | -0.03 | 0.31 | -0.66 | 0.55 | 53.17 |
|  | Slope | -0.08 | 0.39 | -0.84 | 0.68 | 57.69 |
| Habitat | Mosaic | -0.70 | 15.82 | -31.76 | 30.32 | 51.17 |
|  | Mires | -0.03 | 15.82 | -31.09 | 31.01 | 50.17 |
|  | Juniper wood | 0.80 | 15.83 | -30.28 | 31.86 | 52.17 |
|  | Acidic grassland | -0.08 | 15.82 | -31.14 | 30.96 | 50.17 |
| Soil | Peaty gleys | 0.22 | 14.15 | -27.57 | 27.99 | 50.17 |
|  | Organic peaty gleys | 0.37 | 14.15 | -27.42 | 28.13 | 51.17 |
|  | Non-calcareous gleys | -2.65 | 14.19 | -30.52 | 25.19 | 55.18 |
|  | Brown forest Balrownie | 0.66 | 14.15 | -27.13 | 28.43 | 51.17 |
|  | Brown forest Gourdie | 1.40 | 14.16 | -26.40 | 29.18 | 53.17 |
| *With indicator species (DIC = 134.96)* |  |  |  |  |  |  |
|  | *Dryopteris dilatata* | 2.36 | 0.86 | 0.70 | 4.08 | 100.00 |
|  | Soil moisture | 0.57 | 0.32 | -0.04 | 1.20 | 95.21 |
|  | Altitude | -0.81 | 0.45 | -1.72 | 0.05 | 95.19 |
|  | Herbivore damage | -0.65 | 0.37 | -1.41 | 0.05 | 95.20 |
|  | Juniper density | 0.25 | 0.52 | -0.78 | 1.26 | 67.69 |
|  | Intercept | 0.18 | 21.22 | -41.48 | 41.81 | 50.17 |
|  | Berry bearing | 0.06 | 0.26 | -0.45 | 0.57 | 57.68 |
|  | Grazing activity | -0.19 | 0.26 | -0.70 | 0.33 | 75.21 |
|  | Watercourse proximity | -0.18 | 0.33 | -0.85 | 0.45 | 67.70 |
|  | Slope | 0.07 | 0.41 | -0.73 | 0.87 | 55.18 |
| Habitat | Mosaic | -0.97 | 15.82 | -32.03 | 30.06 | 52.18 |
|  | Mires | -0.18 | 15.82 | -31.24 | 30.86 | 50.17 |
|  | Juniper wood | 1.45 | 15.84 | -29.65 | 32.52 | 53.17 |
|  | Acidic grassland | -0.30 | 15.82 | -31.37 | 30.73 | 50.17 |
| Soil | Peaty gleys | -0.14 | 14.15 | -27.93 | 27.63 | 50.17 |
|  | Organic peaty gleys | 0.50 | 14.15 | -27.29 | 28.26 | 51.17 |
|  | Non-calcareous gleys | -3.05 | 14.19 | -30.91 | 24.80 | 57.68 |
|  | Brown forest Balrownie | 1.06 | 14.16 | -26.73 | 28.83 | 52.67 |
|  | Brown forest Gourdie | 1.62 | 14.16 | -26.18 | 29.41 | 54.17 |

The Cairngorms population models all predicted the distribution of symptoms with reasonable accuracy, as the predicted area of symptoms was within 20 % of observed values (Table 4). Addition of a plant indicator improved symptom prediction by 10 % in the Lake District to within 20 % of the observed values. However, predictive model performance was poorer for the Perthshire population where the predicted area of symptomatic juniper was only within 40 % of observed values from both the full subsets and plant indicator models (Table 4). No evidence of overdispersion was found in the residuals of any of the top models (Table 4), nor any consistent, positive, spatial autocorrelation (Appendix D).

The top model produced for the Perthshire population from the full subset selection included all ten possible covariates (DIC 167.45) with only one strong relationship identified between increasing area of symptomatic juniper and increasing soil moisture (Table 5, BCI = 0.70, 4.08). Model fit improved by 32 units (DIC 134.96) when area of *Dryopteris dilatata* was included: a species of large fern that prefers moist, moderately acidic and fertile soils (Table 3). In this model, the strongest effect (BCI did not bridge zero) was increasing area of *P. austrocedri* symptoms with increasing area of *D. dilatata* (BCI = 0.70, 4.08). The area of symptoms also increased very strongly with increasing soil moisture (BCI = -0.04, 1.20) and decreasing altitude (BCI = -1.72, 0.05), and strongly with decreasing area of herbivore damage (BCI = -1.41, 0.05). Six of the seven potential covariates collected across the Lake District population were included in the top model prior to adding indicators (DIC 264.64) with area of *P. austrocedri* symptoms again showing a strong response to soil moisture related covariates, with symptoms strongly increasing with decreasing distance to watercourses (Table 6, BCI = -0.99, 0.08). Including brambles, *Rubus fruticosus* agg., improved the model fit by 56 DIC units (Table 4). The BCI for the relationship between increasing area of symptoms and decreasing distance to watercourses did not bridge zero (BCI = -1.26, -0.14) and the area of symptoms strongly increased with decreasing area of *R. fruticosus* agg. (BCI = -4.19, 0.09), recorded in 12 of 46 quadrats.

**Table 6.** Posterior estimates (mean, standard deviation (SD), 2.5 % and 97.5 % quantiles, and % that does not bridge zero) for fixed effects included in the top model set for the Lake District juniper population.

| Covariate | Mean | SD | 2.5% | 97.5% | % does<br>not bridge<br>zero |
| --- | --- | --- | --- | --- | --- |
| *Without indicator species (DIC = 264.64)* |  |  |  |  |  |
| Intercept | -1.28 | 0.45 | -2.19 | -0.43 | 100.00 |
| Watercourse proximity | -0.41 | 0.27 | -0.99 | 0.09 | 92.70 |
| Soil moisture | 0.09 | 0.25 | -0.41 | 0.58 | 62.69 |
| Juniper density | 0.17 | 0.50 | -0.78 | 1.17 | 62.67 |
| Berry bearing | 0.00 | 0.22 | -0.45 | 0.41 | 52.18 |
| Altitude | 0.22 | 0.24 | -0.24 | 0.70 | 80.19 |
| Slope | -0.26 | 0.28 | -0.83 | 0.29 | 80.20 |
| *With indicator species (DIC = 208.84)* |  |  |  |  |  |
| Intercept | -1.29 | 0.43 | -2.16 | -0.47 | 100.00 |
| Watercourse proximity | -0.66 | 0.29 | -1.26 | -0.14 | 100.00 |
| *Rubus fruticosus agg.* | -1.81 | 1.09 | -4.19 | 0.09 | 95.23 |
| Soil moisture | 0.00 | 0.25 | -0.51 | 0.48 | 50.17 |
| Juniper density | -0.21 | 0.55 | -1.27 | 0.88 | 65.19 |
| Berry bearing | -0.04 | 0.22 | -0.49 | 0.36 | 55.18 |
| Altitude | -0.24 | 0.30 | -0.82 | 0.35 | 77.68 |
| Slope | 0.06 | 0.29 | -0.53 | 0.62 | 57.70 |

Two top models were found for the Cairngorms population including six, and seven, of nine possible covariates; including slope marginally improved model fit (DIC decreased from 226.71 to 225.40). In both models the BCI for soil moisture did not bridge zero, showing a very strong relationship between increasing area of *P. austrocedri* symptoms with increasing soil moisture (Table 7). The individual addition of ten indicator taxa to each of these models resulted in one top model, which contained both slope and cross-leaved heath (*Erica tetralix*). Model fit was improved by 35 and 33 DIC units compared to the full subset selection models (Table 4). Increasing area of symptoms with increasing area of *E. tetralix* was the only strong relationship present, for which the BCI did not bridge zero (BCI = 0.26, 1.28). The only indicator for highly acidic, infertile microsites with high soil moisture is *E. tetralix* recorded in 13 of 50 quadrats (Table 3).

**Table 7.** Posterior estimates (mean, standard deviation (SD), 2.5 % and 97.5 % quantiles, and % that does not bridge zero) for fixed effects included in the top model set for the Cairngorms juniper population.

| Covariate |  | Mean | SD | 2.5% | 97.5% | % does<br>not<br>bridge<br>zero |
| --- | --- | --- | --- | --- | --- | --- |
| *Without indicator species (DIC = 226.71)* |  |  |  |  |  |  |
|  | Soil moisture | 0.54 | 0.22 | 0.09 | 0.98 | 100.00 |
|  | Berry bearing | 0.18 | 0.18 | -0.18 | 0.53 | 82.68 |
|  | Intercept | -1.15 | 27.39 | -54.93 | 52.58 | 51.17 |
|  | Juniper density | -0.18 | 0.44 | -1.01 | 0.70 | 65.20 |
|  | Aspect | -0.02 | 0.21 | -0.45 | 0.39 | 54.17 |
| Habitat | Scots pine woodland | 0.64 | 15.82 | -30.43 | 31.67 | 51.17 |
|  | Heath | 0.19 | 15.82 | -30.86 | 31.22 | 50.17 |
|  | Coniferous plantation | -0.73 | 15.84 | -31.83 | 30.34 | 51.17 |
|  | Acidic grassland | -0.09 | 15.82 | -31.16 | 30.95 | 50.17 |
| Soil | Humus-iron podzols; gleys | 0.46 | 22.36 | -43.44 | 44.33 | 50.17 |
|  | Humus-iron podzols; brown forest | -0.46 | 22.36 | -44.37 | 43.40 | 50.17 |
| *Without indicator species (DIC = 225.40)* |  |  |  |  |  |  |
|  | Soil moisture | 0.57 | 0.23 | 0.12 | 1.01 | 100.00 |
|  | Berry bearing | 0.17 | 0.18 | -0.18 | 0.52 | 82.68 |
|  | Intercept | -1.08 | 27.39 | -54.85 | 52.66 | 51.17 |
|  | Juniper density | -0.24 | 0.44 | -1.09 | 0.65 | 70.19 |
|  | Slope | 0.13 | 0.22 | -0.31 | 0.54 | 72.69 |
|  | Aspect | -0.07 | 0.23 | -0.52 | 0.37 | 60.17 |
| Habitat | Scots pine woodland | 0.67 | 15.82 | -30.39 | 31.71 | 51.17 |
|  | Heath | 0.17 | 15.82 | -30.88 | 31.20 | 50.17 |
|  | Coniferous plantation | -0.76 | 15.84 | -31.86 | 30.31 | 51.17 |
|  | Acidic grassland | -0.07 | 15.82 | -31.14 | 30.97 | 50.17 |
| Soil | Humus-iron podzols; gleys | 0.52 | 22.36 | -43.39 | 44.38 | 50.17 |
|  | Humus-iron podzols; brown forest | -0.52 | 22.36 | -44.42 | 43.35 | 50.17 |
| *With indicator species (DIC = 191.92)* |  |  |  |  |  |  |
|  | *Erica tetralix* | 0.73 | 0.26 | 0.26 | 1.28 | 100.00 |
|  | Soil moisture | 0.19 | 0.27 | -0.36 | 0.71 | 75.21 |
|  | Berry bearing | 0.15 | 0.18 | -0.22 | 0.50 | 77.72 |
|  | Intercept | -1.72 | 27.39 | -55.50 | 52.02 | 52.18 |
|  | Juniper density | -0.02 | 0.46 | -0.88 | 0.91 | 52.68 |
|  | Slope | 0.04 | 0.21 | -0.39 | 0.45 | 57.70 |
|  | Aspect | 0.05 | 0.23 | -0.42 | 0.50 | 57.69 |
| Habitat | Scots pine woodland | 1.12 | 15.82 | -29.95 | 32.16 | 52.67 |
|  | Heath | 0.47 | 15.82 | -30.59 | 31.50 | 51.17 |
|  | Coniferous plantation | -2.26 | 15.85 | -33.39 | 28.84 | 55.18 |
|  | Acidic grassland | 0.69 | 15.83 | -30.39 | 31.73 | 51.17 |
| Soil | Humus-iron podzols; gleys | 0.30 | 22.36 | -43.60 | 44.17 | 50.17 |
|  | Humus-iron podzols; brown forest | -0.30 | 22.36 | -44.21 | 43.56 | 50.17 |

In addition to soil moisture directly measured within quadrats, top models for all populations contained positive effects of juniper density and area of juniper bearing berries and negative effects of slope on symptoms, despite differences in the range of variation sampled across each population (Table 2). Soil and habitat (NVC community) types only included in the Perthshire and Cairngorms models were also always present. None of these covariates showed strong relationships with increasing area of symptoms but removing them resulted in poorer model fit (i.e. the DIC increased by more than two units).

## 4.0 Discussion

Our study provides the first evidence from the northern hemisphere that, out of the wide range of potential abiotic and biotic drivers considered, and despite differences in the range of conditions, geographic location and infection intensity occupied by study populations, soil moisture is the best predictor of *P. austrocedri* symptom distribution in juniper at population scale. This is likely to result from the pathogen’s dependence on soil moisture for zoospore dispersal (Green et al., 2015; Greslebin et al., 2007).

Introductions of non-native *Phytophthora* taxa have been reported from 176 countries across a wide range of climatic zones (Barwell et al., in review). Water availability is commonly identified as an important driver of terrestrial *Phytophthora* distributions at a range of scales. Globally, functional and species diversity increases with precipitation (Redondo et al., 2018). Landscape scale examples include increased incidence of *P. lateralis* in Port Orford cedar with increasing creek drainage area (Jules et al., 2008), and increased *P. ramorum* infection of tanoak with increasing stream proximity (Peterson et al., 2014). In individual trees, the length of *P. cinnamomi* lesions increase in Jarrah with increasing precipitation (Bunny et al., 1995) because water is required to stimulate sporangial formation, trigger zoospore release and enable dispersal (Hardham, 2005).

The importance of soil moisture as a driver of *P. austrocedri* infection in juniper populations is likely to vary between sites with different soil types and hydrology. Area of symptoms increased very strongly with point sampled soil moisture in both the Cairngorms and Perthshire populations, which grow on deep soils formed from glacial till, with pockets of gleying, that retain a high volume of soil moisture throughout the year. This is consistent with population level studies of Chilean cedar where *P. austrocedri* infection increases with soil waterlogging caused by high clay content (La Manna and Rajchenberg, 2004a) or features restricting water permeability (La Manna & Rajchenberg, 2004b).

Though microsite soil moisture is partly a function of soil type, explicitly including soil type in the Cairngorms and Perthshire models always improved model performance but never strongly predicted the area of symptoms, probably because the available data for soil type were too coarse in spatial resolution (250 m) to capture microsite variation.

Spatial variation in area of symptoms was also linked to soil moisture in the Lake District populations but here the strongest association was between symptoms and decreasing proximity to watercourses rather than point sampled soil moisture. Given the steep site topography and freely draining, shallow, sandy soil type, it is likely that juniper in this population is only exposed to long term waterlogging where it grows adjacent to watercourses. Stands of Chilean cedar growing in comparable (freely draining, volcanic) soils also demonstrate increased infection with increasing proximity to watercourses (Calí, 1996; La Manna & Rajchenberg, 2004a).

A key challenge for investigating field scale drivers of disease is obtaining data at a suitably detailed spatial resolution. Modelling microsite soil moisture patterns was prohibited by the availability of fine scale data on hydrological processes (such as precipitation, potential evaporation and runoff generation). Topographic wetness index (TWI), calculated from site topography and watercourse networks, is commonly used as a proxy for soil moisture. In the absence of variability in slope and altitude gradients, the calculation tends to overpredict differences (Grabs et al., 2009) and did not yield an informative distribution map for the Cairngorms juniper population (results not shown). It also assumes uniform soil properties and does not account for complex bedrock surfaces, invalidating the data derived for the Lake District population (results not shown) (Kopecký and Čížková, 2010). Measuring soil moisture directly from stratified quadrats as % volumetric water content captured variation in water table heights but only represents conditions at a single point in time and differences between microsites may be exaggerated by rainfall events that occurred during the sampling period. We introduced the area of vascular plant indicators, selected to represent a range of soil moisture preferences, to test whether such indicators capture longer term water table levels than short term soil moisture field observations, or other fine scale soil attributes affecting transmission and disease such as pH and nitrogen content. This was successful in the Cairngorms, where adding area of cross-leaved heath, *Erica tetralix*, resulted in a very strong, positive, relationship with increasing area of symptoms (Δ DIC = 33), corroborating the response with increasing soil moisture as cross-leaved heath grows in constantly wet but not inundated soils (Hill et al., 2004). These findings suggest microsites most vulnerable to *P. austrocedri* infection could be identified using indicator species with distinctive soil moisture preferences.

Uncoupling relationships between vegetative cover and soil moisture from other factors such as interspecific competition and land management practices proved difficult for the remaining populations. The best Perthshire model was obtained by adding area of broad-buckler fern, *Dryopteris dilatata*, which increased with increasing area of symptoms (Δ DIC = 32, BCI did not bridge zero). This correlation is more likely to result from the fern preferentially colonising dead juniper following *P. austrocedri* induced mortality than suggest a higher percentage of symptoms occurred in the drier soil conditions favoured by the fern (Table 3) (Hill et al., 2004; Rünk et al., 2012).

Adding area of brambles (*Rubus fruticosus* agg.) yielded the greatest improvement in Lake District model performance (Δ DIC = 56) and showed a strong (BCI 0.90 – 0.94), negative relationship with area of symptoms. While four other short-listed taxa for the Lake District also indicate moderate soil moisture conditions, only brambles indicate neutral, moderately fertile soils (Hill et al., 2004). When considered alongside the relationship found in the Cairngorms model between increased symptoms and area of cross-leaved heath, itself an indicator of highly acidic soils (pH 1-3) (Hill et al., 2004), and increasing infection of Chilean cedar in soils containing low levels of alkaline sodium fluoride (NaF) (La Manna et al., 2012) this might suggest *P. austrocedri* occurs more frequently in acidic soils.

However, brambles are also highly unpalatable to herbivores, which preferentially avoid eating them (Bee et al., 2009). Thus increasing symptoms in the absence of brambles might point to herbivore mediated dispersal of inoculum. A cost distance analysis comparing three cattle grazing scenarios (no grazing, roaming with intermittent barriers such as steep slopes and free roaming) found total area and dispersion of *P. austrocedri* was higher in Chilean cedar forests with unrestricted grazing (La Manna et al., 2013). Similarly, infection of Port Orford cedar with *P. lateralis* increases along wildlife (including bear) trails that “fill-in” uninfected sites following disease establishment around creek edges (Jules et al., 2008).

Although our results cannot distinguish between the effects of pH and passive movement of inoculum on herbivore hooves, they do clearly indicate that direct herbivore damage does not increase infection. Herbivore damage was absent from the Cairngorms and Lake District models and though present in Perthshire, symptoms decreased with increasing herbivory (BCI 0.90 – 0.94).

Slope was present in all top models, describing some of the residual variance as a weak, negative relationship with area of symptoms, even in the limited range occupied by the Cairngorms population (Table 2). Positive relationships with area of berry-bearing juniper were also present in all top models and the weak response could indicate female juniper without berries were missed by the survey. Similarly, juniper density was present in all top models showing a weakly positive relationship with symptoms. The intensity of field scale infections of Chilean cedar with *P. austrocedri*, and white oak (*Quercus alba*) with the similarly soil-borne *P. cinnamomi*, were found to increase with increasing host density, suggesting our characterisation of juniper density as ± 20 % cover in 30 × 30 m was too simplistic. This highlights the need for further exploration of the role of host connectivity in facilitating *P. austrocedri* spread in future research across a range of different spatial scales.

Models produced for Perthshire had the lowest accuracy (RMSE 42.59) despite containing the largest number of covariates (11) meaning those included poorly account for the spatial distribution of symptoms. The watercourses mapped for Perthshire include herringbone drainage channels opened in 2011 (Taylor, H., 2019, pers. comm. 1 Nov) after juniper stands started to decline in the late 1990’s (Broome et al., 2008) but before isolation of *P. austrocedri* in 2012 (Green et al., 2015). The drainage work may have inadvertently distributed the pathogen across the site by disturbing watercourses and moving contaminated soil in tyre treads. It is unclear if the very strong increase in area of symptoms with decreasing altitude (BCI > 0.95) reflects the location where the pathogen was first introduced or the drainage activity that was concentrated between 220 – 250 m of the 180 – 310 m altitudinal range (Fig. 3, Table 2) causing the pathogen to spread further and faster than dispersal through soil moisture alone. This highlights the importance of capturing and integrating the spatial arrangement and intensity of management actions into investigations of drivers of site level variation in plant disease impacts (Fernández-habas et al., 2019).

## 5.0 Conclusion

Our study provides valuable insights about how conditions favouring newly invading plant diseases can be delineated by collecting and modelling spatially explicit data at field scale. Directly measuring covariates in the field in sites of different disease status, across key environmental gradients using a carefully designed sampling strategy, enabled us to explore a wide range of potential drivers, identify those with the most explanatory power and make comparisons between populations occupying different ranges along each gradient. In addition, our findings can be used by practitioners to target management interventions that match the inherent scale of pathogen spread.

Interventions to manage juniper populations at local level such as drainage, grazing exclosures and seedling propagation require significant resources (Forestry Commission Scotland, 2006). The introduction of *P. austrocedri* to the UK risks this investment and the longevity of these actions if measures to prevent disease introduction and establishment are not undertaken.

Our results suggest *P. austrocedri* is most likely to infect juniper where it occupies wet microsites - in the UK and across its global range. Surveys by plant health inspectors to detect *P. austrocedri* should be prioritised for populations and stands occupying consistently wet microsites, identification of which could be aided by plant species indicators. While improved biosecurity measures, such as cleaning all machinery, equipment and footwear before and after accessing juniper sites (irrespective of known disease status), will reduce the risk of pathogen introduction and spread (Department for Environment Food & Rural Affairs, 2014b), our results suggest activities that disturb watercourses or wet soils (such as drainage and infrastructure installation) pose the highest risk of collecting and transporting *P. austrocedri* oospores and zoospores.

Our findings further support recommendations published by the UK Department for Environment, Food and Rural Affairs in the Juniper Management Guidelines to divert footpaths away from waterlogged areas and only plant juniper in drier microsites, giving full consideration to the vulnerability of existing populations, suspected disease status, soil type and the watercourse network (Department for Environment Food & Rural Affairs, 2017). Continued emphasis on improving the quality and extent of populations in drier soil conditions by regulating grazing levels, curtailing stand removal and creating spaces for natural regeneration (Broome et al., 2017; Wilkins and Duckworth, 2011) will help maximise resilience of native juniper populations to this new disease threat.

## Supporting information

Appendix A. Mapping juniper study populations

Appendix B. Associate species target list

Appendix C. Distribution of P. austrocedri qPCR results

Appendix D. Additional information for model selection

## Acknowledgements

The authors wish to thank Rothiemurchus Estate, Dalemain Estate, Drummond Estate, the Steeles, and the Taylforths for permitting us to survey juniper populations they manage; Carolyn Riddell for assistance with qPCR analysis, as well as lesion sampling alongside April Armstrong and Ewan Purser; France Gerard and John Redhead for advice on aerial image classification methods; Hollie Cooper, Morag McCracken and Pete Scarlett for preparing and lending survey equipment; and finally to Fiona Cameron, Deborah Comyn-Platt, Etienne Duperron, Rory Hodd, Susan Medcalf, Denise Pallett, Mari Roberts and Rosamund Sparks for their excellent assistance collecting field data. F. Donald was funded by the Scottish Forestry Trust, Scottish Forestry, Forest Research, Scottish Natural Heritage and the Royal Botanic Garden Edinburgh.

## Appendix A. Mapping juniper study populations

## Appendix B. Associate species target list

## Appendix C. Distribution of *P. austrocedri* qPCR results

## Appendix D. Additional information for model selection

