## Appendix A. Mapping juniper study populations for "Small scale variability in soil moisture drives infection of vulnerable juniper populations by invasive forest pathogen"

To identify sampling locations along a gradient of juniper density, juniper distribution maps were first required for each population.

**A.1.1 Distribution mapping of the Perthshire juniper population**

A map of the Perthshire population was created from a classification of imagery collected and processed by Caledonian Air Surveys Ltd (Whittome, 2010) under contract to Scottish Natural Heritage (SNH). Full colour (RGB) and false colour infrared (CIR) imagery was collected in August 2010 at 15 cm resolution from a light aircraft using a 50-megapixel Hasselblad H4D-50 digital camera (Whittome, 2010). Maximum likelihood (ML) classification of the resulting imagery was implemented in the Focus program included in Geomatica version 9.1.7-R5 (PCI Geomatics, 2010). The classification used five object classes: asymptomatic juniper, dead juniper, trees, bracken and other. The spectral signature space occupied by each object class is assigned a multivariate Gaussian distribution, with a mean and covariance matrix fit for each class from sets of training pixels (Richards, 1999). Each pixel in the image is then assigned by the ML classifier to the object class with the highest likelihood (Environmental Systems Research Institute, 2016). No information about the number of training pixels used for classification or the resulting classification accuracy was supplied (Whittome, 2010). Two shapefiles of juniper, classified as asymptomatic or dead, were produced and made available to the authors by SNH. We joined the shapefiles in R version 3.4.0 (R Core Team, 2017) and converted it to a raster, resampled to 1 m resolution (i.e. the minimum unit of juniper increased from 0.15 m^2^ to 1 m^2^) using the *rasterize* function in the raster package (Hijmans, 2016) implemented with a mean function. The raster was then overlain with a 10 x 10 m grid created from 5 m digital elevation models (DEM) supplied by NeXTPerspectives™ (updated February 2014), averaged to 10 m using the *aggregate* function in the raster package (Hijmans, 2016).

**A.1.2 Distribution mapping of the Lake District and Cairngorms juniper populations**

No pre-existing spatial information was available for the Lake District or Cairngorms juniper populations, so we used ARC GIS v. 10.5 (Environmental Systems Research Institute, 2017) to carry out a supervised ML classification of full colour (RGB), 25 cm imagery collected in 2010 and supplied by NeXTPerspectives™. One image was provided for each location, which was aligned and clipped to the boundary of the corresponding juniper population then classified using the ML method described above. A dataset of pixels representing different observable object classes was created for each image and partitioned into a training set and a test set at a ratio of 80:20 pixels (Table A.1). As few *P. austrocedri* symptoms were visible across either population in 2010, separation of asymptomatic and symptomatic juniper object classes was not required for classification.

Table A.1. List of object classes, number of training and test pixels used to classify 25 cm RGB images of the Lake District and Cairngorms juniper populations.

| **Population** | **Object classes** | **Training** | **Test** |
| --- | --- | --- | --- |
| Lake District | Juniper | 54000 | 10800 |
|  | Trees | 47271 | 9454 |
|  | Tree shadow | 496 | 99 |
|  | Rock | 1514 | 303 |
|  | Other | 156124 | 31225 |
| Cairngorms | Juniper | 35057 | 7011 |
|  | Trees | 121016 | 24203 |
|  | Tree shadow | 6643 | 1329 |
|  | Rock | 6060 | 1212 |
|  | Bracken | 5881 | 1176 |
|  | Grass | 56072 | 11214 |
|  | Heath | 62335 | 12467 |

Classification was performed using the *create signatures* and *maximum likelihood classification* tools in the spatial analyst toolbox. The value of each pixel in the test dataset was extracted from the classified image using *extract* in the raster package (Hijmans, 2016) implemented in R. version 3.4.0 (R Core Team, 2017). Confusion matrices were used to summarise the probability of correctly assigning test pixels to object classes (Table A.2). Commission is the number of pixels incorrectly included in an object class whereas omission is the number of pixels incorrectly missed out; user’s accuracy is 1 - commission error and producer’s accuracy is 1 - omission error. Cohen’s kappa statistic was calculated using the *accuracy* function in the rfUtilities package (Evans et al., 2011) as a commonly used metric to interpret test pixel classification accuracy, accounting for both commission and omission errors and correcting overall prediction accuracy by the accuracy expected to occur by chance (Allouche et al., 2006). The following indicative kappa thresholds were suggested by Landis and Koch (1977): 0.41 – 0.6 moderate, 0.61 – 0.8 substantial and > 0.81 almost perfect.

As the kappa statistics for juniper classification met the threshold for “moderate” and “substantial” classification accuracy in the Cairngorms and Lake District populations respectively, the analysis was considered sufficiently accurate at the time to identify potential sampling locations described in the quadrat stratification section in the methods.

The resulting rasters of juniper distribution were then resampled to 1 m resolution and overlain with a 10 x 10 m grid using the same methods described for the Perthshire population.

Table A.2. Confusion matrix presenting rates of omission and commission, user’s and producer’s accuracies per object class obtained from the ML classification of the 25 cm RGB imagery supplied by Next Perspectives™ of a) the Lake District b) the Cairngorms juniper study populations.

|  |  | **Reference** | | | | | Total | User's | Commission | Kappa |
| --- | --- | --- | --- | --- | --- | --- | --- | --- | --- | --- |
|  |  | Juniper | Trees | Tree shadow | Rock | Other |  |  |  |  |
| **Predicted** | Juniper | 7655 | 441 | 35 | 70 | 2969 | 11170 | 0.69 | 0.31 | 0.62 |
|  | Trees | 758 | 8388 | 14 | 0 | 847 | 10007 | 0.84 | 0.16 | 0.83 |
|  | Tree shadow | 260 | 219 | 49 | 3 | 244 | 775 | 0.06 | 0.94 | 0.11 |
|  | Rock | 543 | 1 | 1 | 190 | 6198 | 6933 | 0.03 | 0.97 | 0.04 |
|  | Other | 1584 | 405 | 0 | 40 | 20967 | 22996 | 0.91 | 0.09 | 0.54 |
|  | Total | 10800 | 9454 | 99 | 303 | 31225 |  |  |  |  |
|  | Producer's | 0.71 | 0.89 | 0.49 | 0.63 | 0.67 |  |  |  |  |
|  | Omission | 0.29 | 0.11 | 0.51 | 0.37 | 0.33 |  |  |  |  |

a)

|  |  | **Reference** | | | | | | | Total | User's | Commission | Kappa |
| --- | --- | --- | --- | --- | --- | --- | --- | --- | --- | --- | --- | --- |
|  |  | Juniper | Trees | Tree shadow | Rock | Bracken | Grass | Heath |  |  |  |  |
| **Predicted** | Juniper | 4483 | 2489 | 0 | 2 | 0 | 469 | 2712 | 10155 | 0.44 | 0.56 | 0.44 |
|  | Trees | 2210 | 20981 | 11 | 0 | 0 | 785 | 0 | 23987 | 0.87 | 0.13 | 0.80 |
|  | Tree shadow | 0 | 604 | 1311 | 0 | 0 | 0 | 0 | 1915 | 0.68 | 0.32 | 0.99 |
|  | Rock | 0 | 0 | 0 | 1197 | 7 | 0 | 10 | 1214 | 0.99 | 0.01 | 0.81 |
|  | Bracken | 0 | 0 | 0 | 1 | 1149 | 0 | 0 | 1150 | 1.00 | 0.00 | 0.99 |
|  | Grass | 17 | 120 | 0 | 3 | 11 | 9601 | 1 | 9753 | 0.98 | 0.02 | 0.90 |
|  | Heath | 301 | 9 | 7 | 9 | 9 | 359 | 9744 | 10438 | 0.93 | 0.07 | 0.44 |
|  | Total | 7011 | 24203 | 1329 | 1212 | 1176 | 11214 | 12467 |  |  |  |  |
|  | Producer's | 0.64 | 0.87 | 0.99 | 0.99 | 0.98 | 0.86 | 0.78 |  |  |  |  |
|  | Omission | 0.36 | 0.13 | 0.01 | 0.01 | 0.02 | 0.14 | 0.22 |  |  |  |  |

b)

**A.1.3 Revised distribution mapping of the Lake District juniper population**

Concerned that inaccuracy in the image classification may lead to errors in proportional sampling of the eight juniper density categories, we wanted to compare the density distribution of sampled quadrats to those derived from the original, and an improved, image classification. The quality of the Cairngorms image was insufficient to make re-analysis practicable but the opportunity arose to re-classify the Lake District image using ARC GIS v.10.5 after fieldwork was completed. We addressed the uneven illumination by separating the image, by eye, into five different sections with different depths of shadow and developing sets of 70 training to 30 test pixels unique to each section (Table A.3). Multiple studies have shown maximum likelihood classification of land cover is outperformed by using the random forest (RF) algorithm (Attarchi and Gloaguen, 2014; Khatami et al., 2016; Waske and Braun, 2009). We implemented the algorithm using the *random trees* tool in the spatial analyst toolbox. Maximum number of trees was set to 200, maximum tree depth to 50 and maximum number of samples per class to 1000. The resulting ESRI classification definition file was then loaded in the *classify raster* tool in the spatial analyst toolbox to produce classified rasters of each image section, assessed using confusion matrices and kappa as outlined in S.1.2. Once the best classification of juniper according to the accuracy statistics (minimum kappa of 0.7) was obtained for each section, the classified image sections were mosaiced back together for the remaining processing.

Predicted juniper pixels isolated from any other juniper pixels in eight neighbouring directions were identified and deleted using the *freq* function in the raster package (Hijmans, 2019) implemented in R v. 3.5.2 (R Core Team, 2018). Juniper pixels visible in the centre of deciduous tree crowns were then manually deleted. These operations deleted 0.05 % juniper pixels identified by the classifier.

Table A.3. List of object classes and total number of pixels used to train the RF algorithm and test the result of the classification of 25 cm RGB image of the Lake District juniper population.

| **Class** | **Training** | **Test** |
| --- | --- | --- |
| Juniper | 17747 | 7606 |
| Trees | 36782 | 15764 |
| Tree shadow | 9369 | 4041 |
| Rock | 15231 | 6528 |
| Bracken | 49218 | 21092 |
| Grass | 9541 | 4158 |
| Heath | 1828 | 784 |

Compared to the original Lake District image analysis, the revised classification of juniper improved by 0.13 kappa units keeping it within the range for substantial classification accuracy (Table A.4). Where the original classification predicted 29 % more juniper pixels than the number observed, the revised classification predicted 15 % fewer juniper pixels (Table A.2a, Table A.4). User’s accuracy improved to 82 % reducing the number of pixels incorrectly classified as juniper by 13 % and producer’s accuracy improved to 74 % increasing the number of pixels correctly classified as juniper by 3 % (Table A.2a, Table A.4). The largest proportion of pixels mistakenly identified as juniper were donated from the “tree shadow” category (10 %) and the greatest percentage of misidentified juniper pixels were predicted to be grass (9 %).

Table A.4. Confusion matrix presenting rates of omission and commission, user’s and producer’s accuracies per object class obtained from the RF revised classification of the 25 cm RGB image supplied by Next Perspectives™ of the Lake District juniper population.

|  |  | **Reference** | | | | | | | Total | User's | Commission | Kappa |
| --- | --- | --- | --- | --- | --- | --- | --- | --- | --- | --- | --- | --- |
|  |  | Juniper | Trees | Tree shadow | Rock | Bracken | Grass | Heath |  |  |  |  |
| **Predicted** | Juniper | 5626 | 396 | 668 | 80 | 61 | 42 | 17 | 6890 | 0.82 | 0.18 | 0.75 |
|  | Trees | 503 | 13474 | 100 | 4 | 987 | 344 | 0 | 15412 | 0.87 | 0.13 | 0.82 |
|  | Tree shadow | 275 | 116 | 3187 | 12 | 12 | 0 | 14 | 3616 | 0.88 | 0.12 | 0.82 |
|  | Rock | 57 | 43 | 50 | 5641 | 1183 | 34 | 6 | 7014 | 0.8 | 0.2 | 0.82 |
|  | Bracken | 303 | 52 | 5 | 605 | 17537 | 478 | 5 | 18985 | 0.92 | 0.08 | 0.81 |
|  | Grass | 667 | 1660 | 19 | 34 | 1305 | 3251 | 0 | 6936 | 0.47 | 0.53 | 0.55 |
|  | Heath | 175 | 23 | 17 | 12 | 8 | 0 | 742 | 977 | 0.76 | 0.24 | 0.84 |
|  | Total | 7606 | 15764 | 4046 | 6388 | 21093 | 4149 | 784 |  |  |  |  |
|  | Producer's | 0.74 | 0.85 | 0.79 | 0.88 | 0.83 | 0.78 | 0.95 |  |  |  |  |
|  | Omission | 0.26 | 0.15 | 0.21 | 0.12 | 0.17 | 0.22 | 0.05 |  |  |  |  |

Pixels classified as juniper in the revised image were subset to a new raster, then resampled to 1 m using *rasterize* in the raster package (Hijmans, 2016) implemented with a mean function. The same method outlined in the quadrat stratification section was used to calculate the area of juniper at 10 x 10 and 30 x 30 m scales for each cell and then assign it to one of the eight juniper density categories (Table A.5). The ML analysis identified 10795 10 x 10 m cells containing juniper, compared to 8399 in RF classification. The proportion of cells distributed across density categories were comparable between the original and revised classifications (Figure A.1, Table A.5). Few cells were found in categories with dense 10 x 10 m but sparse 30 x 30 m cover (categories 3-4), or sparse 10 x 10 m and dense 30 x 30 m cover (category 5). The greatest proportion of cells were allocated to categories 7-8 with high % cover at both scales but the category with the highest proportion of cells differed by method. The sum of the area of juniper calculated at 10 x 10 m and 30 x 30 m for each juniper cell is highly correlated (Pearson r^2^ = 0.69) between the two classifications. Comparing the sample distribution to that of the revised image classification suggests too many quadrats were sampled at high juniper density (category 8) but this may arise from the slight under-prediction of juniper occurrence in the revised image classification (Table A.4). The sample distribution does match the revised classification well in the lower density categories (Figure A.1), so we are satisfied that juniper sampling based on the original image classification was roughly proportional to the juniper density actually present across the site.

Table A.5. Description of the eight juniper density categories based on % juniper cover in 10 x 10 m quadrats and the 30 x 30 m including each quadrat, the percentage of cells allocated to each category following classification of 1 m resolution RGB imagery supplied by Next Perspectives™ using ML (original) and RF (revised) algorithms and the percentage of cells sampled by the field survey in each category.

| Density category | % juniper cover | | % 10 x 10 m cells per category | | |
| --- | --- | --- | --- | --- | --- |
|  | 30 x 30 m | 10 x 10 m | Original | Revised | Sample |
| 1 | 0 - 20 | 1 – 10 | 9 | 14 | 11 |
| 2 |  | 11 – 25 | 5 | 13 | 11 |
| 3 |  | 26 – 50 | 1 | 2 | 0 |
| 4 |  | 50 - 100 | 0 | 0 | 0 |
| 5 | 21 - 100 | 1 – 10 | 3 | 4 | 0 |
| 6 |  | 11 – 25 | 12 | 15 | 13 |
| 7 |  | 26 – 50 | 28 | 29 | 26 |
| 8 |  | 50 - 100 | 42 | 23 | 39 |

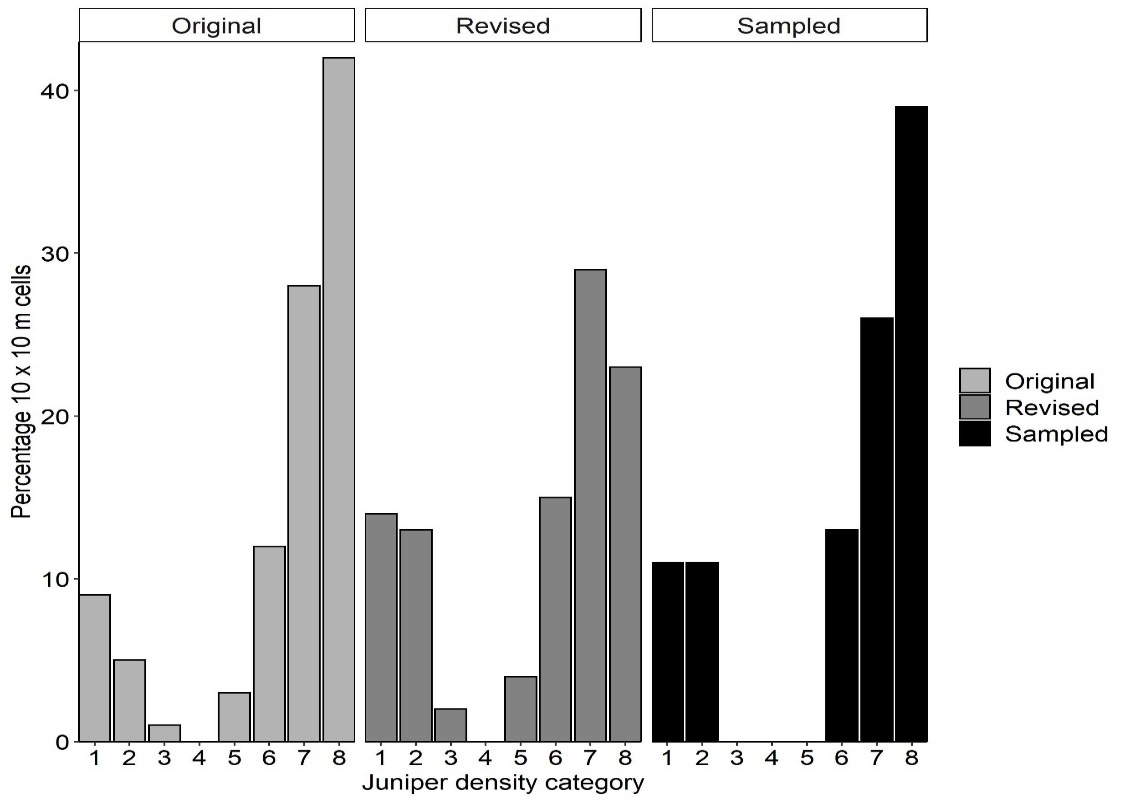

Figure A.1. Distribution of 10 x 10 m cells in the Lake District study area across eight juniper density categories (Table A.5) comparing the classification of 1 m resolution RGB imagery supplied by Next Perspectives™ using the original (ML) and revised (RF) methods and the percentage of cells sampled by the field survey. The number of 10 x 10 m cells containing juniper compared is 10795 (original), 8399 (revised), 46 (sampled).

Environmental Systems Research Institute, 2017. ArcGIS Desktop, version 10.5. Redlands, CA.

Environmental Systems Research Institute, 2016. How maximum likelihood classification works [WWW Document]. URL <http://desktop.arcgis.com/en/arcmap/10.3/tools/spatial-analyst-toolbox/how-maximum-likelihood-classification-works.htm>

Evans, J.S., Murphy, M.A., Holden, Z.A., Cushman, S.A., 2011. Modeling species distribution and change using Random Forests, in: Drew, C.A., Wiersma, Y.F., Huettmann, F. (Ed.), Predictive Species and Habitat Modeling in Landscape Ecology: Concepts and Applications. New York, pp. 139–159.

Hijmans, R.J., 2016. raster: geographic data analysis and modeling. R package version 3.0-2. <https://CRAN.R-project.org/package=raster>

Khatami, R., Mountrakis, G., Stehman, S.V., 2016. A meta-analysis of remote sensing research on supervised pixel-based land-cover image classification processes: general guidelines for practitioners and future research. Remote Sens. Environ. 177, 89–100. <https://doi.org/10.1016/j.rse.2016.02.028>

Landis, J.R., Koch, G.G., 1977. The measurement of observer agreement for categorical data. Biometrics 33, 159–174. <https://doi.org/10.2307/2529310>

PCI Geomatics, 2010. Geomatica version 9.1.7-R5. Markham, Canada.

R Core Team, 2018. R: A language and environment for statistical computing. R Foundation for Statistical Computing. Vienna, Austria. URL <https://www.R-project.org/>.

R Core Team, 2017. R: A language and environment for statistical computing. R Foundation for Statistical Computing. Vienna, Austria. URL <https://www.R-project.org/>.

Richards, J.A., 1999. Remote sensing and digital image analysis: an introduction, 2nd ed. Springer Berlin, Heidelberg.

Waske, B., Braun, M., 2009. Classifier ensembles for land cover mapping using multitemporal SAR imagery. ISPRS J. Photogramm. Remote Sens. 64, 450–457. <https://doi.org/10.1016/j.isprsjprs.2009.01.003>

Whittome, T., 2010. Report No CASL-0909-129-1a Glenartney juniper wood aerial photography 2010. Report commissioned by Scottish Natural Heritage, Inverness.
