## Appendix B. Associate species target list for "Small scale variability in soil moisture drives infection of vulnerable juniper populations by invasive forest pathogen"

**Additional information for model selection**

**D.1 Correlations between environmental covariates**

Correlations between all covariates investigated in the Perthshire, Lake District and Cairngorms juniper populations were investigated (Figure D.1). The dependent variable “Symptoms” (area of symptomatic juniper in 10 x 10 m) and the independent variable “Area” (area of juniper in 10 x 10 m) were included in the analysis but collinearity was only examined between the covariates (Figure D.1). No covariates were correlated with a Pearson r^2^ value ≥ 0.6.

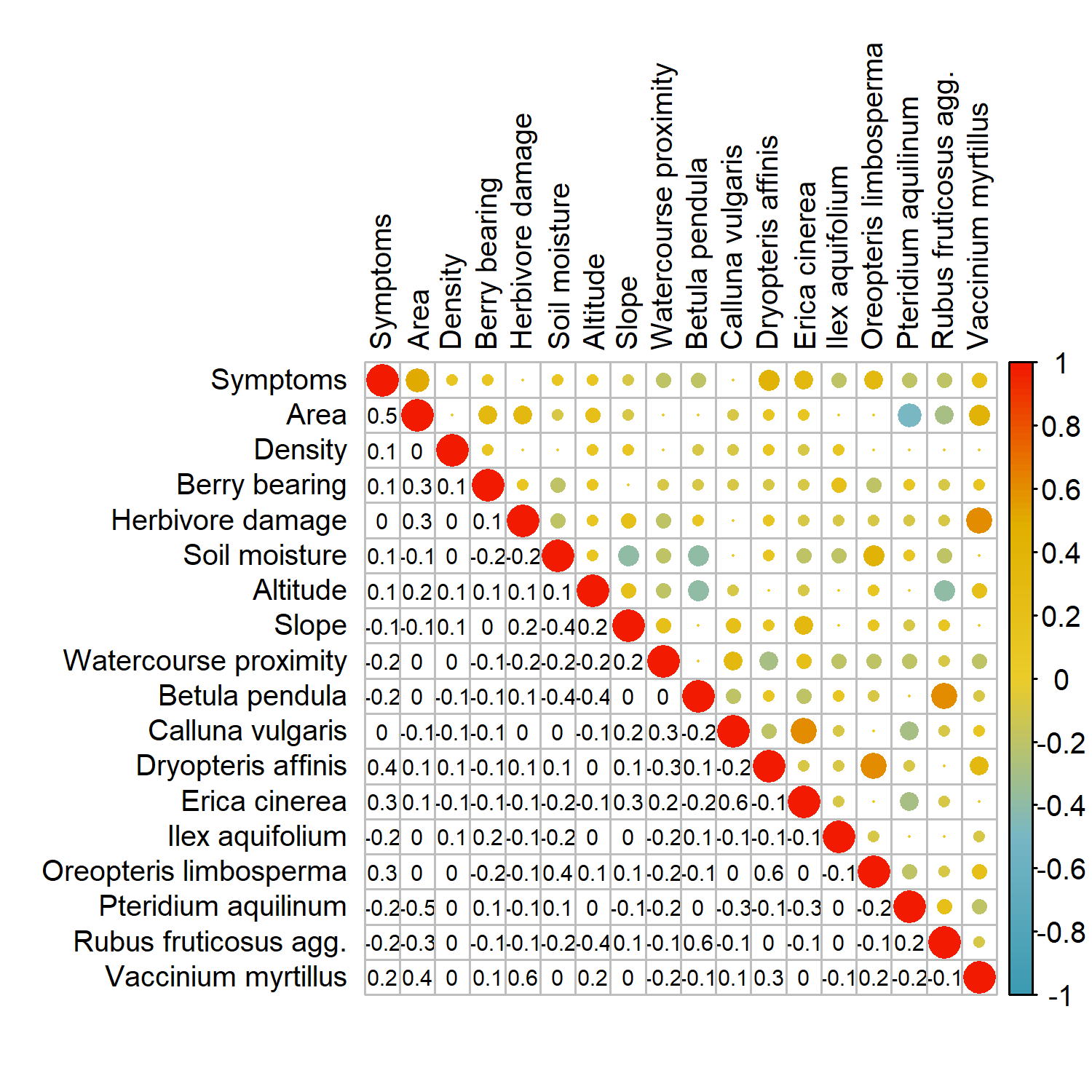

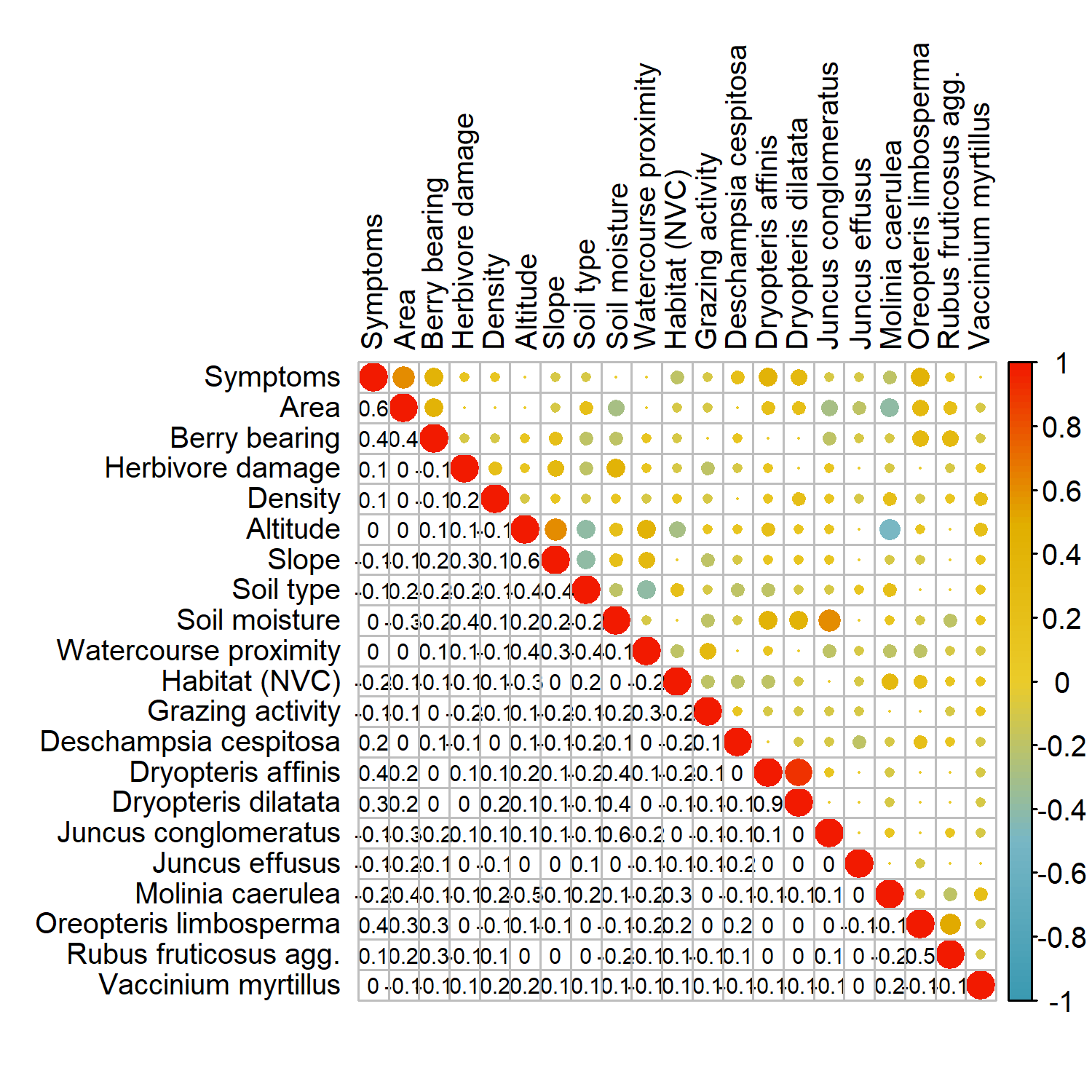

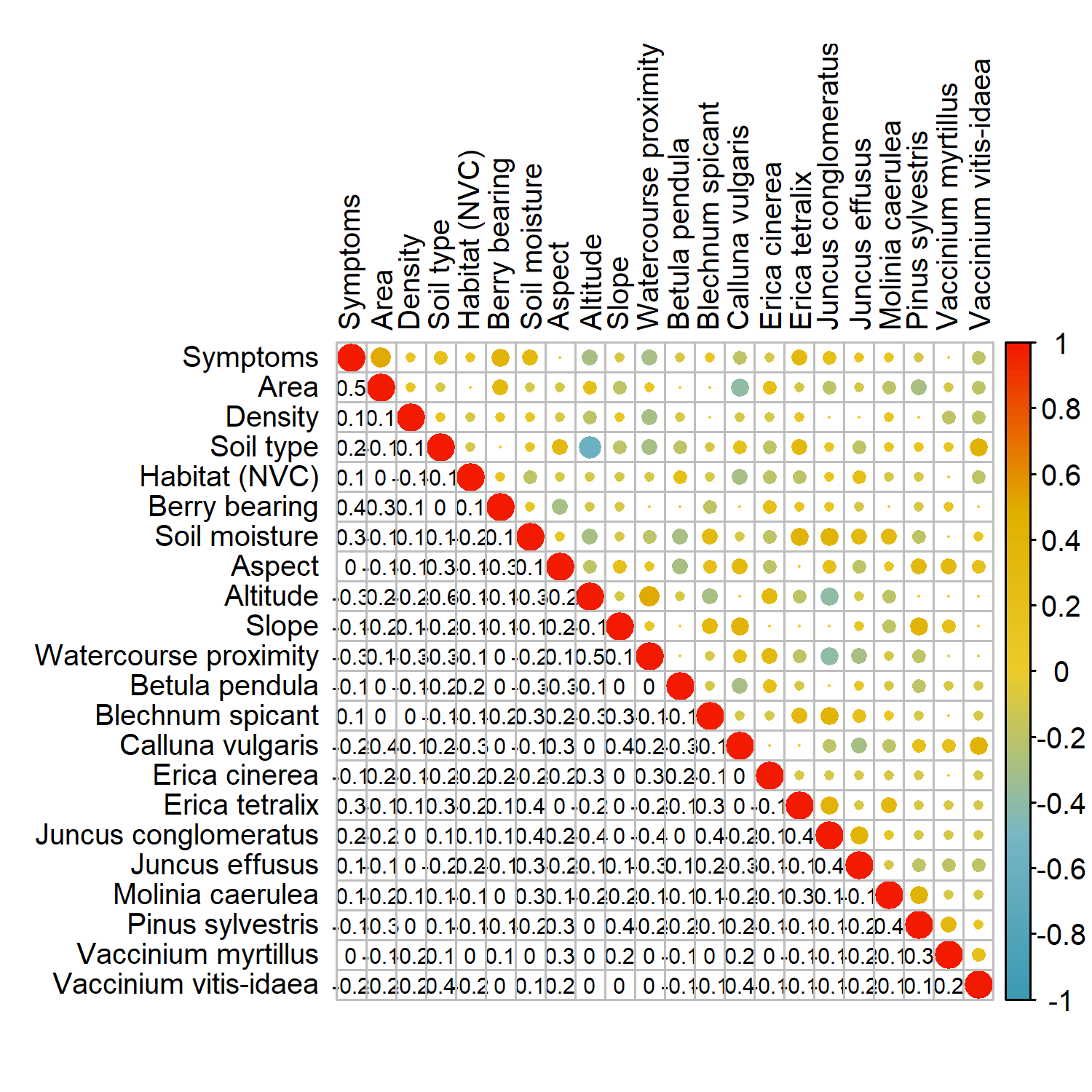

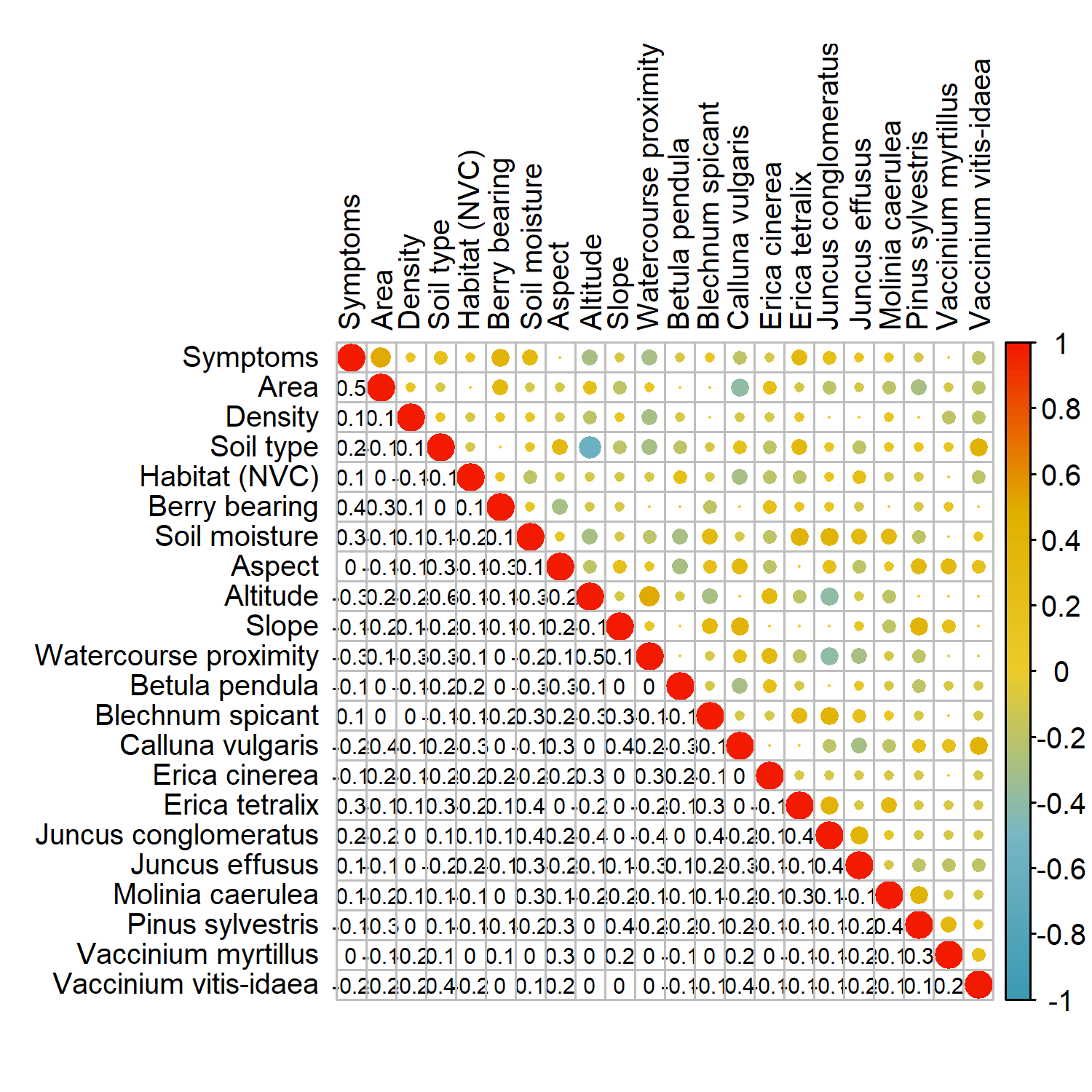

Perthshire	Lake District	Cairngorms

Figure D.1: Correlation plot between all investigated metrics for the Perthshire, Lake District and Cairngorms juniper populations. Pearson r^2^ values are shown using colour scale and text to 1 decimal place. Covariate descriptions and units of measurement are given on Table 1.

**D.2. Accounting for spatial autocorrelation in model residuals**

Correlations in the residuals generated from beta-binomial GLMMs were calculated using Moran’s I statistic. Positive spatial autocorrelation of the residuals was rarely significant in any of the top models for any of the populations up to the first 1000 m of inter-quadrat distance (Figure D.2).

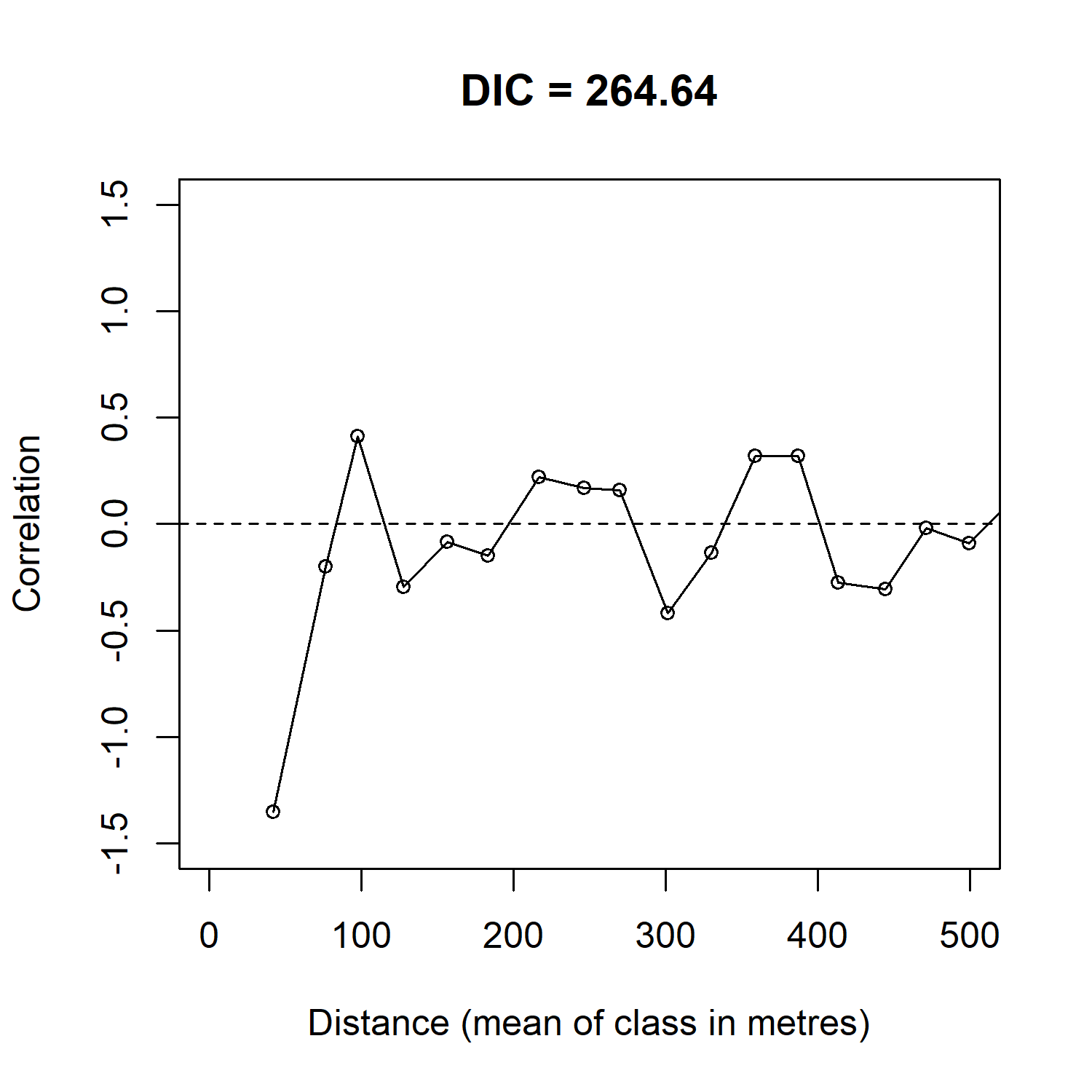

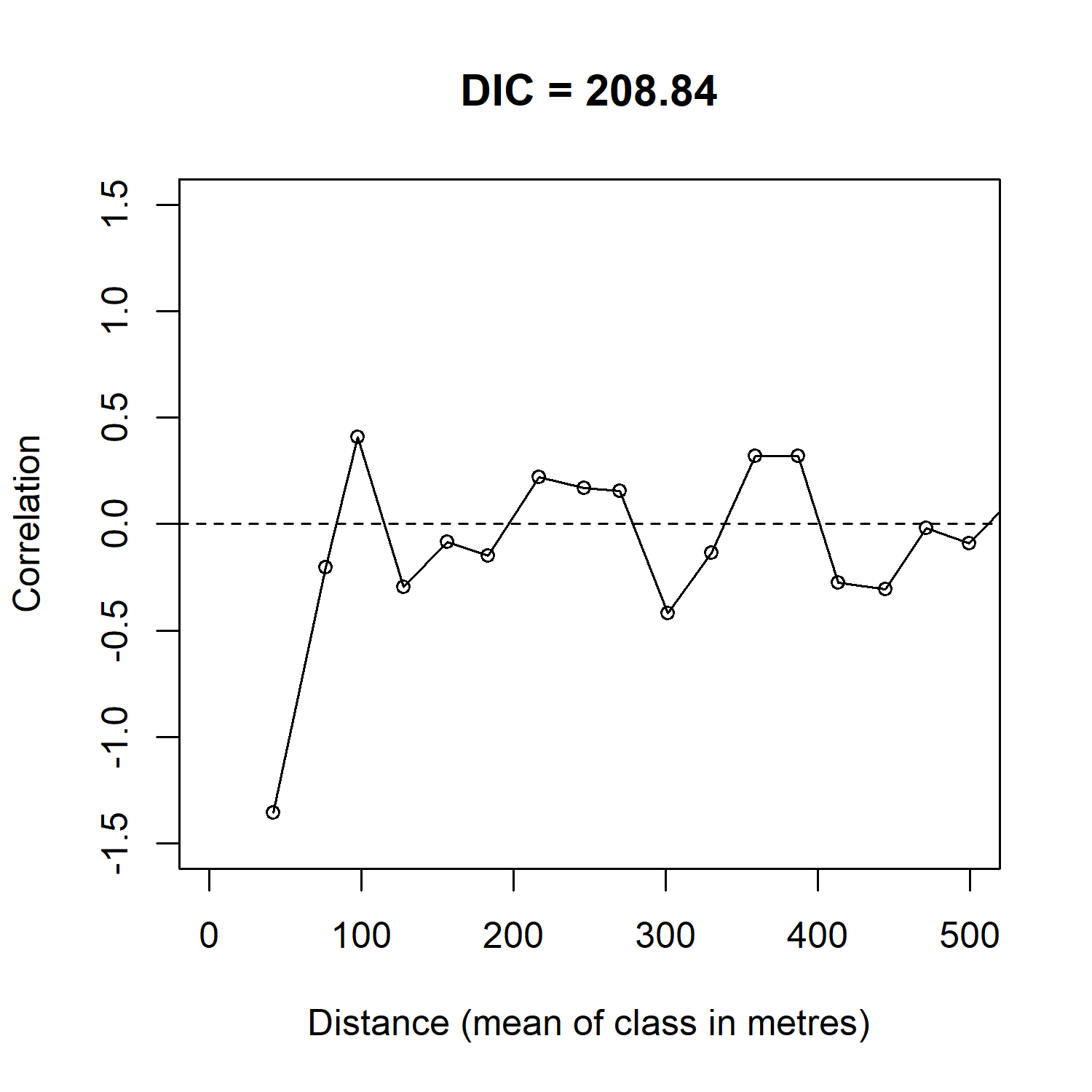

Lake District

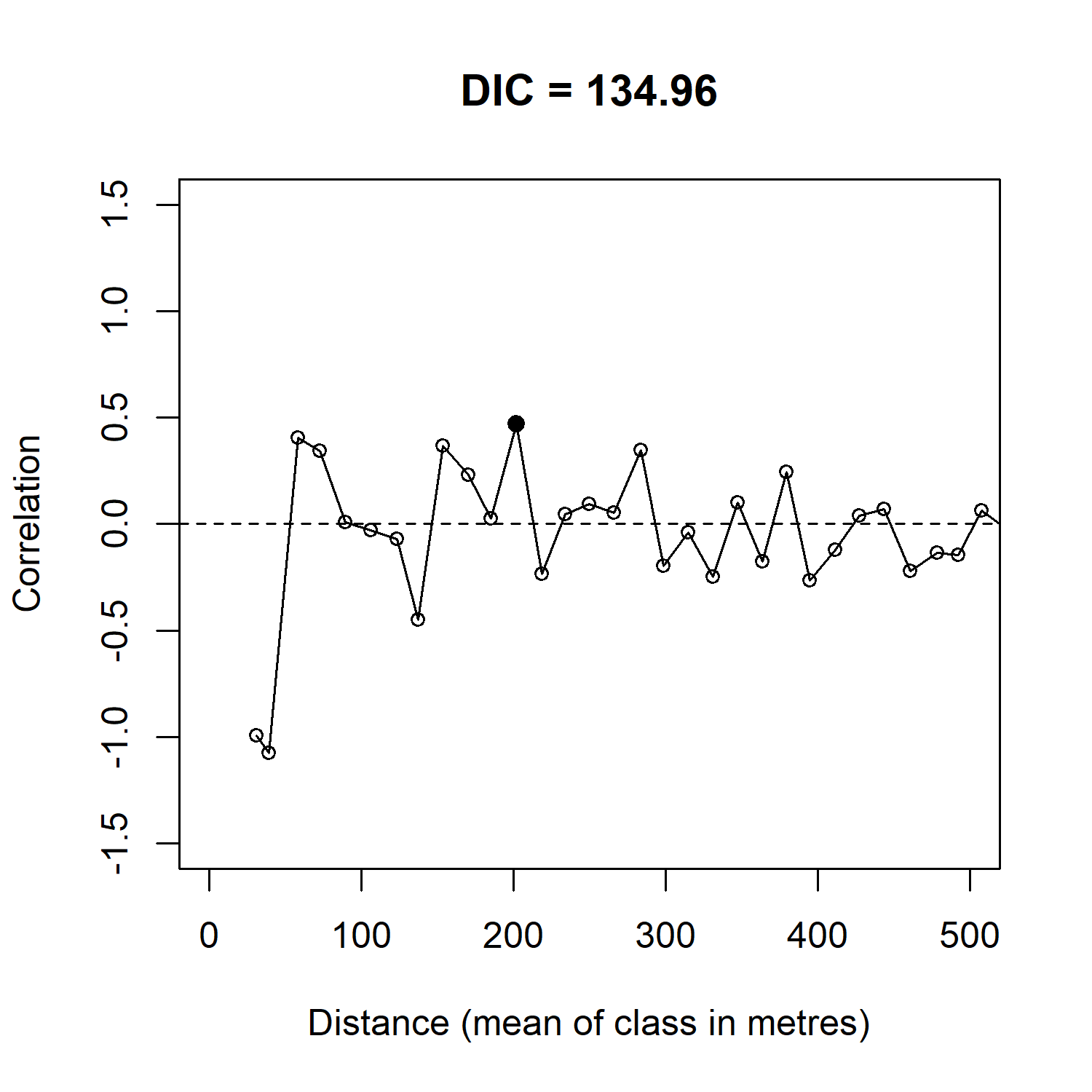

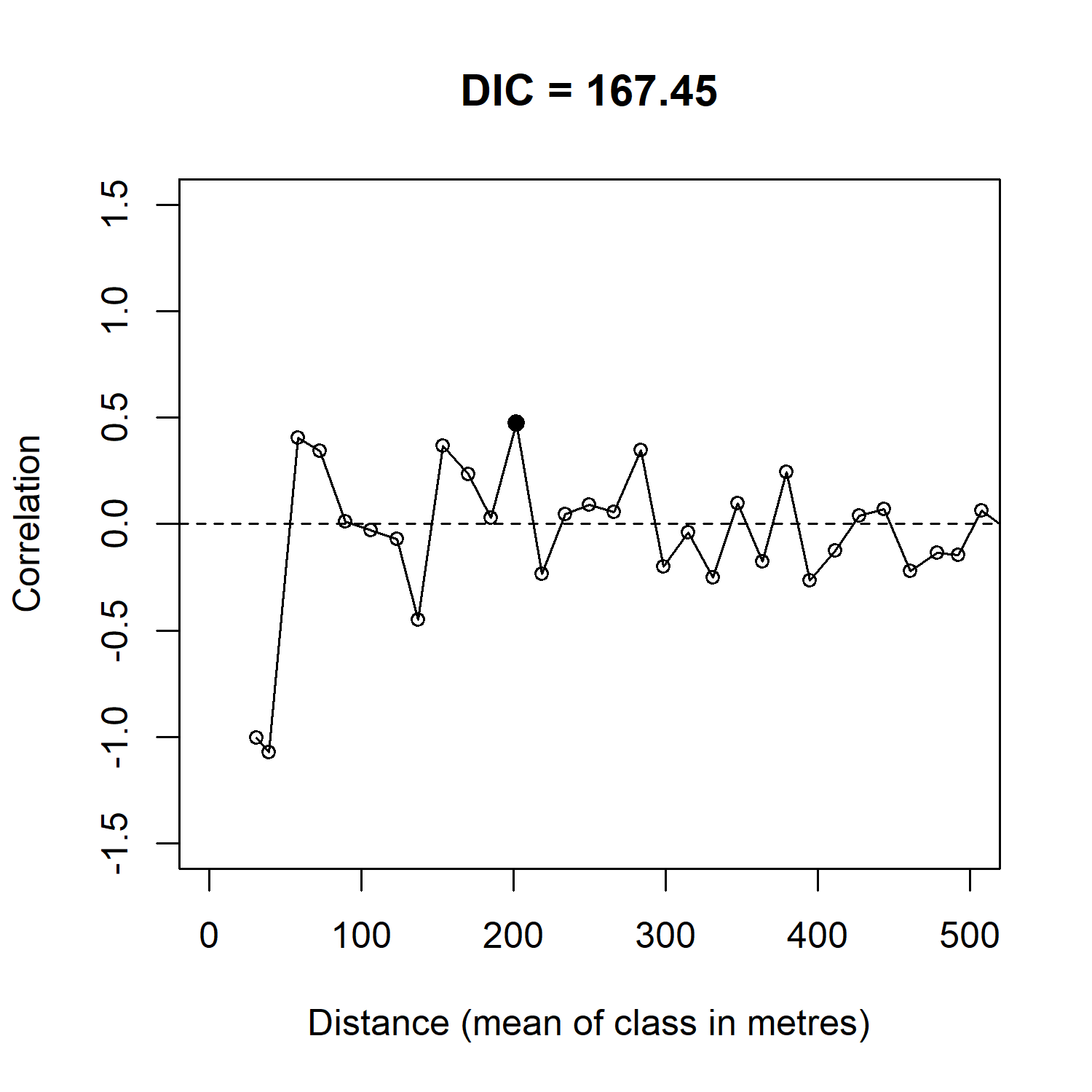

Perthshire

Cairngorms

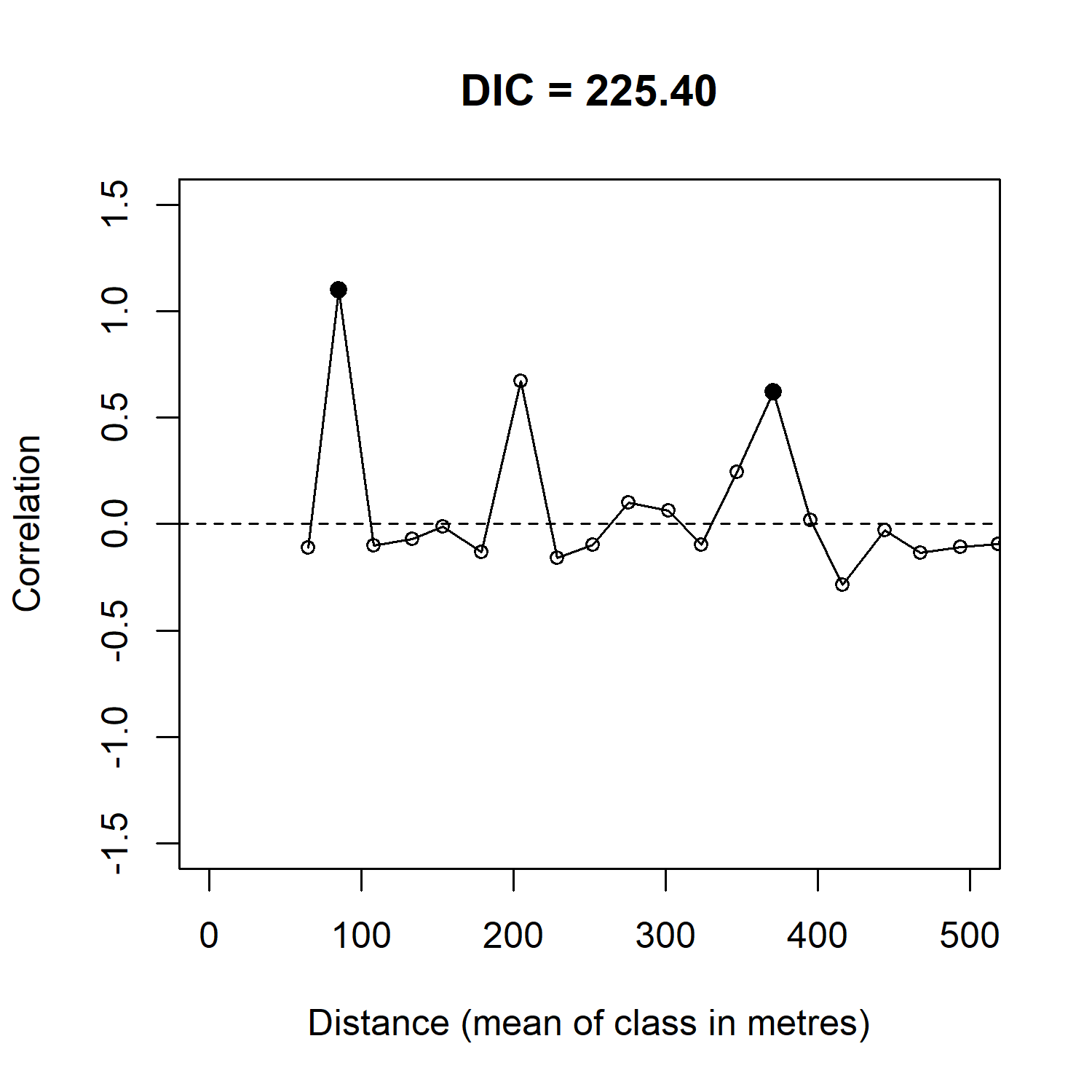

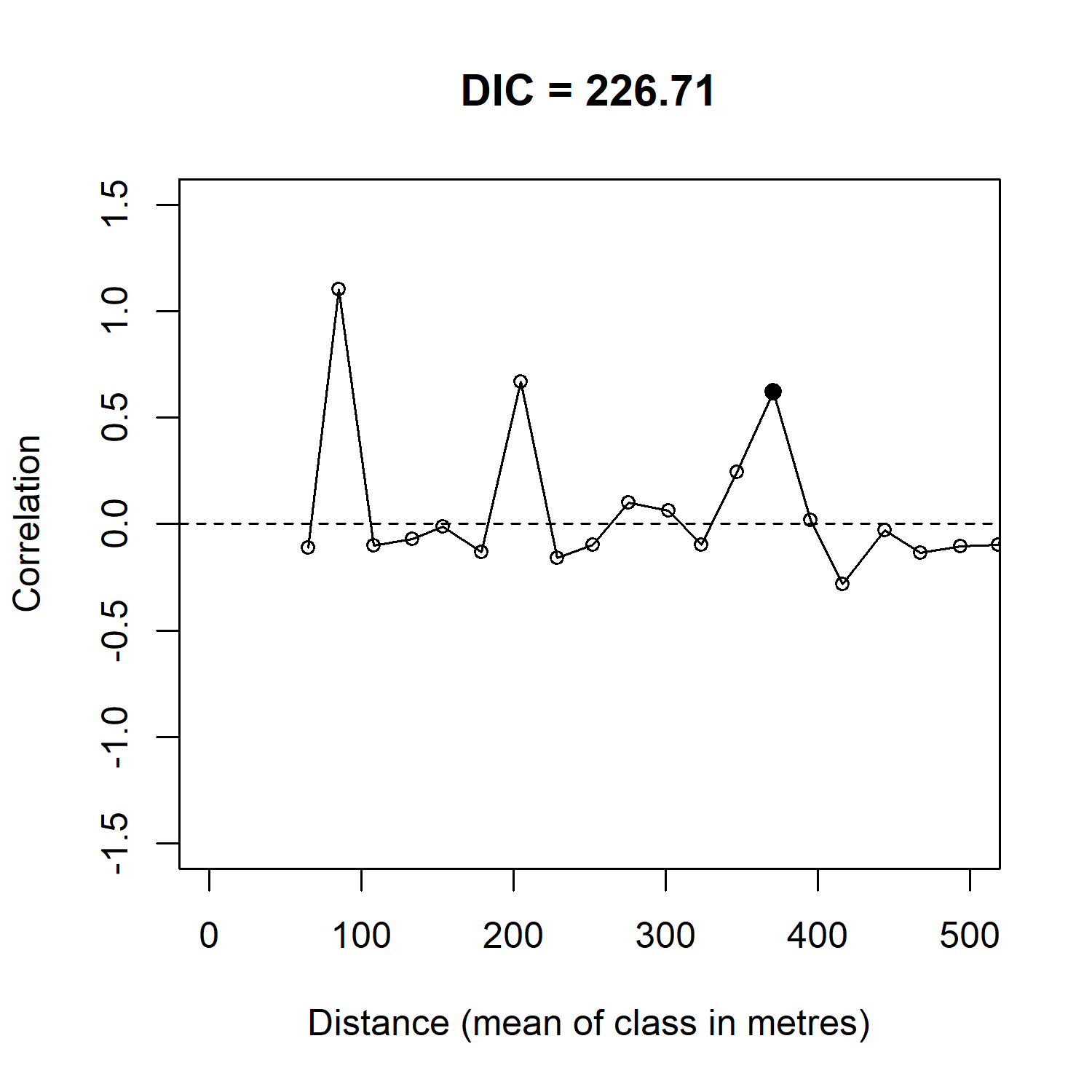

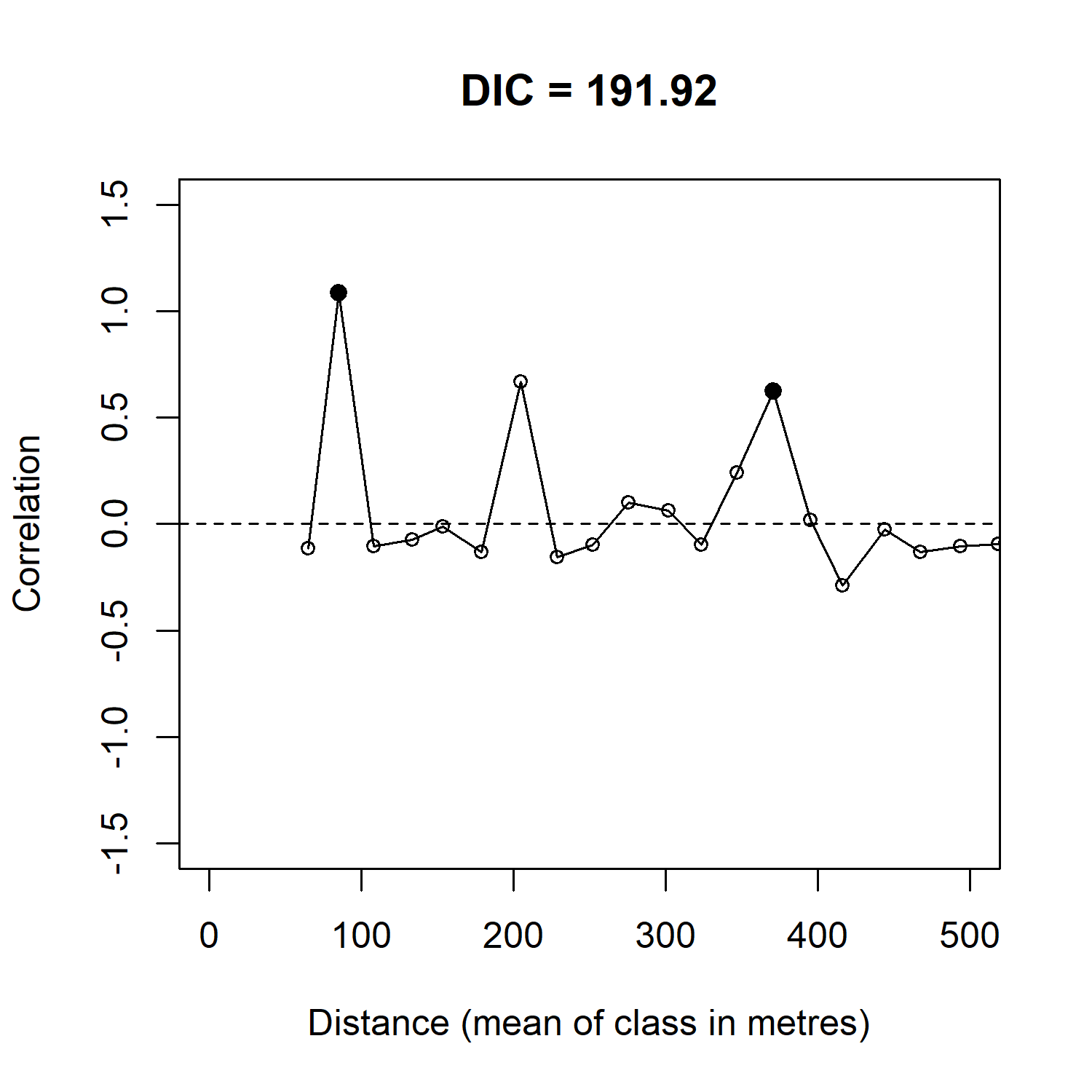

Figure D.2:

Moran’s I p-values calculated from the residuals of the top set of beta-binomial GLMs plotted against the inter-cell distance (m). Significant p-values (95 % confidence interval) are shown in black.
