## Appendix C. Distribution of P. austrocedri qPCR results for "Small scale variability in soil moisture drives infection of vulnerable juniper populations by invasive forest pathogen"

Tissue from lesions symptomatic for *P. austrocedri* was collected from juniper with foliage symtpoms within field quadrats and tested for *P. austrocedri* DNA using qPCR. Positive results were obtained for 49 % of quadrats in Perthshire, 58 % in the Lake District and 60 % in the Cairngorms (Table C.1), distributed across the full extent of each juniper population (Figure C.1). Symptomatic lesions were not found in the remaining quadrats, with the exception of three quadrats (5 %) in Perthshire and three (6 %) in the Lake District, where the presence of DNA was not positively confirmed by qPCR (Table C.1, Figure C.1). Inability to detect lesions on dead trees explains the inverse relationship between the percentage of quadrats with positive qPCR results and the area of symptoms detected across each study population (Figure 3, Table 2).

Table C.1**.** Comparison of the numbers of quadrats, lesion samples, positive and “not detected” qPCR results for *P. austrocedri* DNA for each study population. No detection of *P. austrocedri* occurs where symptomatic lesions could not be found within quadrats or where qPCR did not confirm DNA presence.

| Measurement | Perthshire | Lake District | Cairngorms |
| --- | --- | --- | --- |
| Number of 10 x 10 m quadrats surveyed | 51 | 48 | 50 |
| Number of lesion samples collected | 28 | 31 | 30 |
| Number of quadrats where *P. austrocedri* present | 25 | 28 | 30 |
| Number of quadrats where *P. austrocedri* not detected | 26 (3) | 20 (3) | 20 (0) |

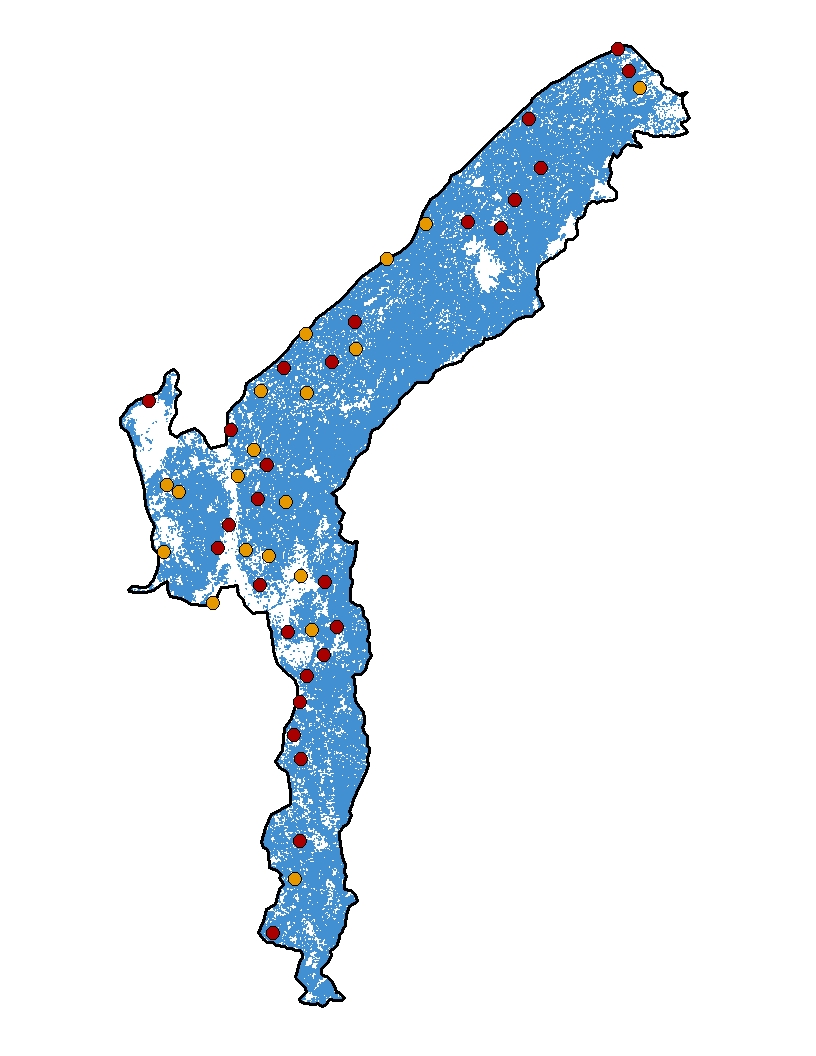

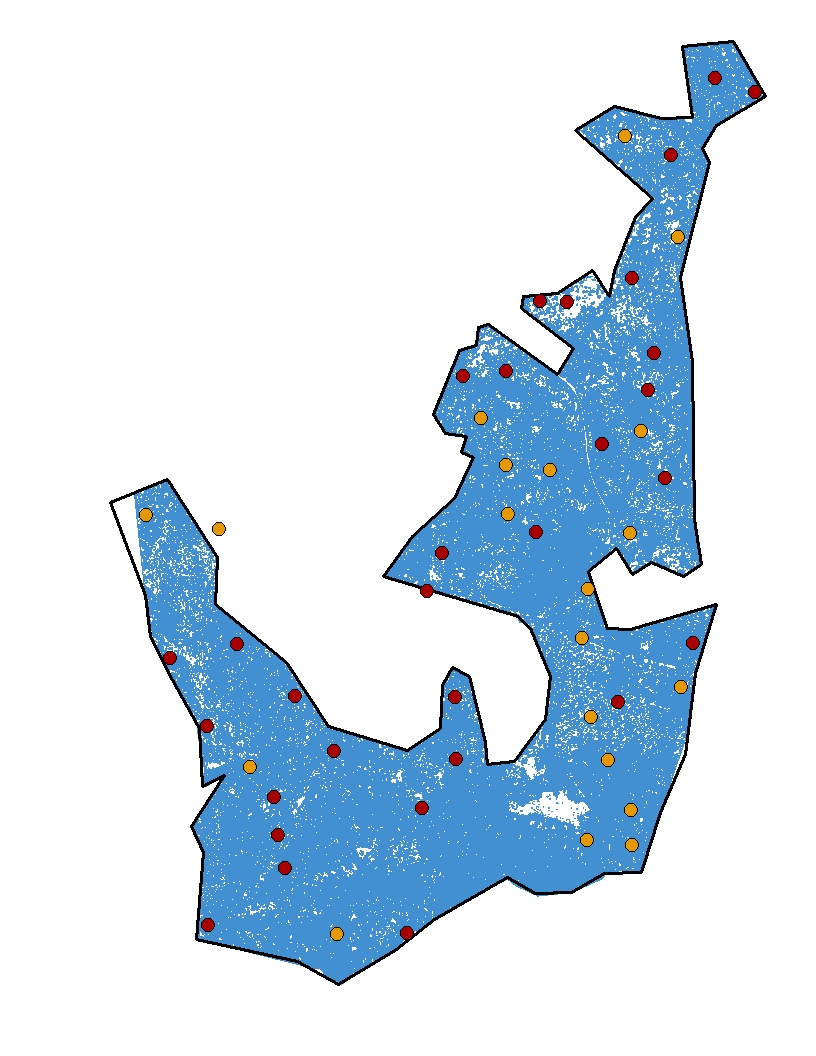

**400 m**

**400 m**

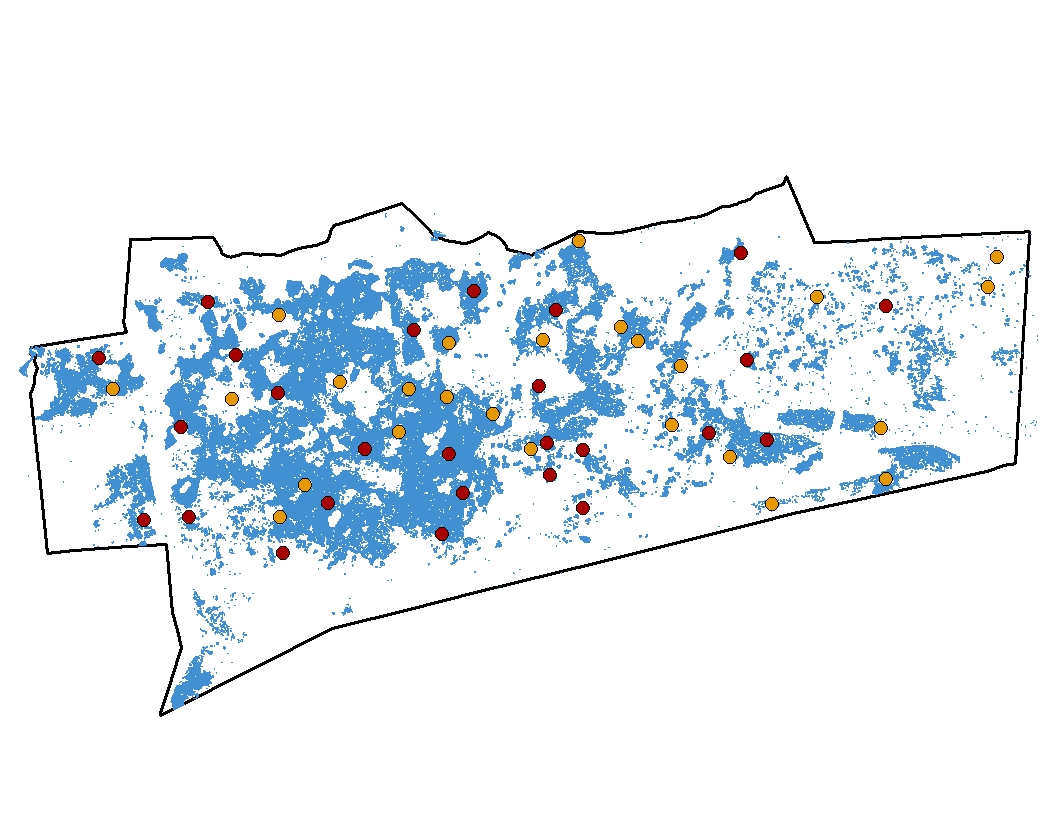

**400 m**

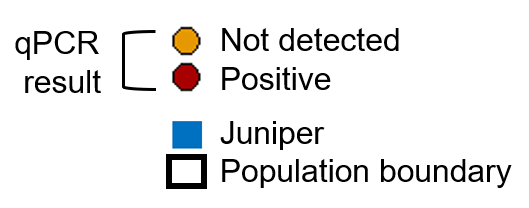

**Cairngorms**

**Lake District**

**Perthshire**

Figure C.1: Spatial distribution of qPCR results obtained from lesions symptomatic for *P. austrocedri* collected from 10 x 10 m quadrats at each study population.
