## Appendix D. Additional information for model selection for "Small scale variability in soil moisture drives infection of vulnerable juniper populations by invasive forest pathogen"

**Associate species target list**

Table B.1. List of 42 target vascular plant species to record in 10 x 10 m quadrats categorised using Ellenberg moisture (F), reaction (R) and nitrogen (N) values given in Hill, Preston, & Roy (2004)**.**

| Vascular plant taxon | Soil pH & fertility | | | Soil moisture | | |
| --- | --- | --- | --- | --- | --- | --- |
|  | Highly acidic  (R = 2), nutrient poor  (N = 1-2) | Slightly acidic  (R = 3-5), moderately fertile  (N = 3-5) | Neutral  (R = 6-7), fertile  (N = 6) | High moisture (F = 8-9) | Moderate moisture  (F = 6-7) | Lower moisture  (F = 5) |
| *Erica tetralix* | *x* |  |  | *x* |  |  |
| *Calluna vulgaris* | *x* |  |  |  | *x* |  |
| *Empetrum nigrum* | *x* |  |  |  | *x* |  |
| *Pinus sylvestris* | *x* |  |  |  | *x* |  |
| *Vaccinium myrtillus* | *x* |  |  |  | *x* |  |
| *Arctostaphylos uva-ursi* | *x* |  |  |  |  | *x* |
| *Erica cinerea* | *x* |  |  |  |  | *x* |
| *Vaccinium vitis-idaea* | *x* |  |  |  |  | *x* |
| *Betula pendula* |  | *x* |  |  |  | *x* |
| *Fagus sylvatica* |  | *x* |  |  |  | *x* |
| *Ilex aquifolium* |  | *x* |  |  |  | *x* |
| *Luzula sylvatica* |  | *x* |  |  |  | *x* |
| *Pteridium aquilinum* |  | *x* |  |  |  | *x* |
| *Quercus robur* |  | *x* |  |  |  | *x* |
| *Rubus fruticosus agg.* |  | *x* |  |  |  | *x* |
| *Taxus baccata* |  | *x* |  |  |  | *x* |
| *Ulex europaeus* |  | *x* |  |  |  | *x* |
| *Alnus glutinosa* |  |  | *x* | *x* |  |  |
| *Iris pseudacorus* |  |  | *x* | *x* |  |  |
| *Fraxinus excelsior* |  |  | *x* |  | *x* |  |
| *Acer pseudoplatanus* |  |  | *x* |  |  | *x* |
| *Coryllus avellana* |  |  | *x* |  |  | *x* |
| *Crataegus monogyna* |  |  | *x* |  |  | *x* |
| *Hedera helix* |  |  | *x* |  |  | *x* |
| *Ulmus glabra* |  |  | *x* |  |  | *x* |
| *Molinia caerulea* |  |  |  | *x* |  |  |
| *Myrica gale* |  |  |  | *x* |  |  |
| *Salix cinerea* |  |  |  | *x* |  |  |
| *Athyrium filix-femina* |  |  |  |  | *x* |  |
| *Betula pubescens* |  |  |  |  | *x* |  |
| *Deschampsia cespitosa* |  |  |  |  | *x* |  |
| *Dryopteris affinis* |  |  |  |  | *x* |  |
| *Dryopteris dilitata* |  |  |  |  | *x* |  |
| *Dryopteris filix-mas* |  |  |  |  | *x* |  |
| *Juncus conglomeratus* |  |  |  |  | *x* |  |
| *Juncus effusus* |  |  |  |  | *x* |  |
| *Lonicera periclymenum* |  |  |  |  | *x* |  |
| *Oreopteris limbosperma* |  |  |  |  | *x* |  |
| *Quercus petraea* |  |  |  |  | *x* |  |
| *Sorbus aucuparia* |  |  |  |  | *x* |  |
